# High-resolution mapping of DNA alkylation damage and base excision repair at yeast transcription factor binding sites

**DOI:** 10.1101/2021.09.24.461700

**Authors:** Mingrui Duan, Smitha Sivapragasam, Jacob S. Antony, Jenna Ulibarri, John M. Hinz, Gregory M.K. Poon, John J. Wyrick, Peng Mao

## Abstract

DNA base damage arises frequently in living cells and needs to be removed by base excision repair (BER) to prevent mutagenesis and genome instability. Both the formation and repair of base damage occur in chromatin and are conceivably affected by DNA-binding proteins such as transcription factors (TFs). However, to what extent TF binding affects base damage distribution and BER in cells is unclear. Here, we used a genome-wide damage mapping method, *N*-methylpurine-sequencing (NMP-seq), to characterize alkylation damage distribution and BER at TF binding sites in yeast cells treated with the alkylating agent methyl methanesulfonate (MMS). Our data shows that alkylation damage formation was mainly suppressed at the binding sites of yeast TFs Abf1 and Reb1, but individual hotspots with elevated damage levels were also found. Additionally, Abf1 and Reb1 binding strongly inhibits BER *in vivo* and *in vitro*, causing slow repair both within the core motif and its adjacent DNA. The observed effects are caused by the TF-DNA interaction, because damage formation and BER can be restored by depletion of Abf1 or Reb1 protein from the nucleus. Thus, our data reveal that TF binding significantly modulates alkylation base damage formation and inhibits repair by the BER pathway. The interplay between base damage formation and BER may play an important role in affecting mutation frequency in gene regulatory regions.

## Introduction

DNA in living cells is exposed to an array of genotoxic agents, both endogenous and exogenous. Alkylating agents comprise a large number of reactive chemicals present in cells and in the environment (Fu et al., 2012), which can react with the nitrogen and oxygen atoms of DNA bases to induce formation of alkylation damage. Some alkylation damage is cytotoxic and mutagenic (Kondo et al., 2010), and thus poses threats to cell growth and genome stability. On the other hand, the cytotoxicity of DNA alkylation is utilized in chemotherapy. Alkylating agents such as temozolomide (TMZ) are used for the treatment of glioblastoma and other cancers (Fu et al., 2012; Newlands et al., 1997). Therefore, studies of alkylation damage and its repair are relevant for both cancer prevention and therapy.

The most common alkylation lesions are *N*-methylpurines (NMPs), including 7-methylguanine (7meG) and, to a lesser extent, 3-methyladenine (3meA) (Kondo et al., 2010). Although 7meG is not genotoxic by itself, it is prone to spontaneous depurination to form a mutagenic apurinic (AP) site (Fu et al., 2012). 7meG can also form deleterious DNA-protein crosslinks with the lysine-rich histone tails (Yang et al., 2018). The 3meA damage is even more harmful than 7meG, as 3meA lesions block DNA polymerases and affect DNA replication (Plosky et al., 2008). Hence, NMP lesions need to be repaired in a timely manner to avoid detrimental outcomes such as cell death or mutations. The primary repair pathway for NMPs is base excision repair (BER), which is initiated by alkyladenine-DNA glycosylase (AAG; also known as MPG and ANPG) in human cells or its yeast ortholog Mag1 (Wyatt et al., 1999). During BER, AAG/Mag1 removes the alkylated base and generates an AP site, which is then cleaved by the apurinic/apyrimidinic endonuclease (APE1) (Whitaker and Freudenthal, 2018). Subsequently, DNA polymerase and ligase are recruited to the nick to conduct repair synthesis and ligation, respectively (Krokan and Bjørås, 2013).

Transcription factors (TFs) are key proteins that regulate gene expression. Many TFs bind to DNA in a sequence-specific manner to direct transcription initiation to target promoters (Jolma et al., 2013). While TFs mainly function in transcriptional regulation, their binding to DNA can affect DNA damage formation and repair (Mao and Wyrick, 2019). To this end, several TF proteins have been shown to modulate formation of ultraviolet (UV) light-induced photolesions (Frigola et al., 2021; Hu et al., 2017; Mao et al., 2018) and inhibit nucleotide excision repair (NER) (Conconi et al., 1999; Sabarinathan et al., 2016). The altered UV damage formation and suppressed NER are believed to cause increased mutation rates at TF binding sites in skin cancers (Frigola et al., 2021; Mao et al., 2018; Sabarinathan et al., 2016). Previous studies have also found that mutation rates are significantly increased at TF binding sites in non-UV exposed tumors (Kaiser et al., 2016; Melton et al., 2015), such as gastric and prostate cancers (Guo et al., 2018; Morova et al., 2020). However, what causes the high mutation rates in non-UV exposed cancers remains elusive. Since base damage (e.g., oxidative, alkylation, uracil, and so on) caused by endogenous and exogenous damaging sources is prevalently associated with cancer mutations (Tubbs and Nussenzweig, 2017; Wallace et al., 2012), a potential mechanism for mutation elevation in non-UV exposed tumors is increased base damage formation and/or suppressed BER in TF-bound DNA. However, this hypothesis has not been tested and it is unclear to what extent TF binding affects base damage formation and BER.

Alkylation damage has been widely used as a model lesion for BER studies (Fu et al., 2012; Li et al., 2015). We previously developed an alkylation damage mapping method, *N*-methylpurine-sequencing (NMP-seq), to precisely map 7meG and 3meA lesions in cells treated with methyl methanesulfonate (MMS) (Mao et al., 2017). Here, we used NMP-seq to analyze alkylation damage formation and BER at the binding sites of ARS binding factor 1 (Abf1) and rDNA enhancer binding protein 1 (Reb1), two essential yeast TFs that have been extensively characterized. The genome-wide binding sites for Abf1 and Reb1 have been identified at near base-pair resolution (Kasinathan et al., 2014; Rossi et al., 2021) and the DNA-binding mechanisms were analyzed in previous studies (Jaiswal et al., 2016; McBroom and Sadowski, 1994a). Analysis of our NMP-seq data indicates that both damage formation and BER are affected by TF binding in yeast cells. We further show that Reb1 protein binding directly inhibits BER of alkylation damage *in vitro*. Collectively, these analyses uncover an important role for TF binding in modulating base damage formation and inhibiting BER.

## Results

### Abf1 and Reb1 modulate alkylation damage formation at their binding sites

NMP-seq is a sequencing method developed to map 7meG and 3meA lesions across the genome (Mao et al., 2017). This method employs BER enzymes AAG and APE1 to digest MMS-damaged DNA and create a nick at the NMP lesion site, which is then ligated to adaptor DNA for next-generation sequencing (Supplemental Fig. S1A). As NMP lesion sites are precisely tagged by the adaptor DNA, sequencing with a primer complementary to the adaptor generates a genome-wide profile of NMP lesions at single-nucleotide resolution (Mao et al., 2017).

To determine how TF binding affects NMP lesion formation, we analyzed initial NMP lesions at Abf1 and Reb1 binding sites in yeast immediately after 10 min MMS treatment (i.e., no repair incubation). The ongoing BER during the period of MMS exposure may repair some of the damage and affect analysis of NMP formation. To minimize the effect of endogenous BER, we used a BER-deficient *mag1* deletion strain (i.e., *mag1*Δ) to profile the initial NMP distribution. We obtained a total of ~44 million sequencing reads in MMS-treated *mag1*Δ cells. The majority of the reads (~56%) were associated with G nucleotides (G reads), followed by A nucleotides (A reads) (Supplemental Fig. S1B), consistent with the expected trend of 7meG and 3meA lesion formation after MMS treatment (Friedberg et al., 2006).

Since 7meG is the major class of lesion induced by MMS, we first characterized 7meG formation at Abf1 and Reb1 binding sites. To account for potential DNA sequence bias at the binding sites, we also mapped NMP damage in naked yeast genomic DNA, in which all proteins were removed and the purified DNA was damaged by incubating with MMS (Supplemental Fig. S1C and S1D). Normalization of cellular G reads by the naked DNA G reads enables us to elucidate the modulation of 7meG formation by TF proteins. Importantly, we found that formation of 7meG was significantly inhibited at Abf1 and Reb1 binding sites relative to the flanking DNA (Fig. 1A and 1B, Supplemental Fig. S2A). Analysis of the average 7meG levels in 5 bp non-overlapping moving windows indicates that 7meG was reduced by up to 40% and 70% for Abf1 and Reb1 binding sites, respectively. Furthermore, the extent of damage reduction was correlated with the level of TF occupancy, as Reb1 binding sites with low occupancy (occupancy <10) (Kasinathan et al., 2014) only slightly reduced 7meG formation (Fig. 1C).

**Figure 1.**
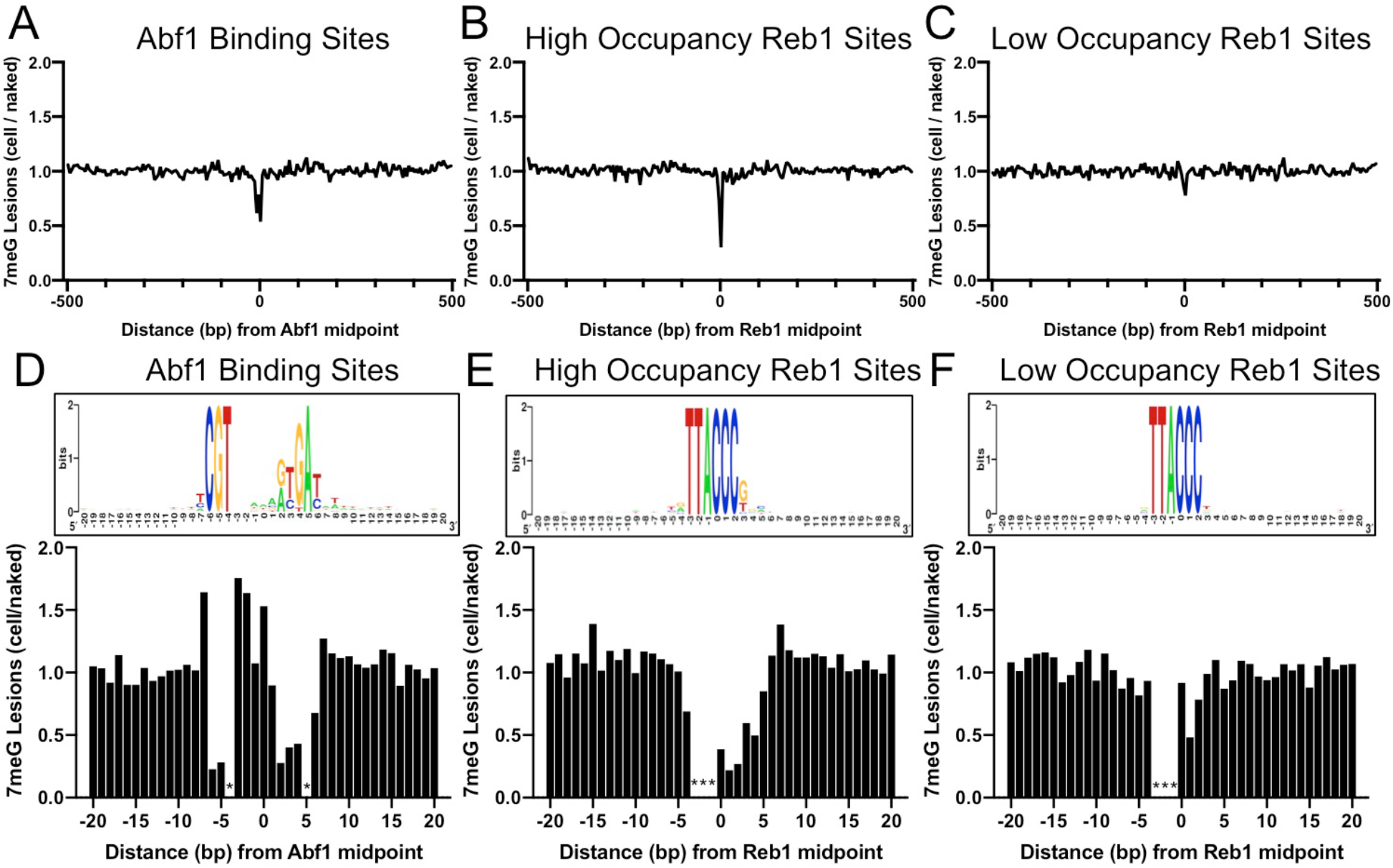
Formation of 7meG lesions at Abf1 and Reb1 binding sites. **(A)** Distribution of 7meG damage at 661 Abf1 binding sites and the flanking DNA in MMS-treated yeast cells. The cellular (i.e., *mag1*Δ-0 h) 7meG levels in 5 bp non-overlapping moving windows were normalized to damage in naked yeast DNA. The normalized ratio was scaled to 1.0 and plotted along the aligned Abf1 sites. **(B)** Distribution of 7meG at 784 ‘high-occupancy’ Reb1 binding sites and the flanking DNA. NMP-seq data was analyzed at Reb1 binding sites. **(C)** Distribution of 7meG at 472 ‘low-occupancy’ Reb1 binding sites. **(D)** to **(F)** High-resolution plots showing 7meG formation in the Abf1, ‘high-occupancy’, and ‘low-occupancy’ Reb1 binding motif and the immediately adjacent DNA. The top panel depicts the consensus motif sequence for each transcription factor. The lower panel shows the normalized damage levels and each column points to a specific position at the binding site. Asterisks indicate conserved motif positions with exclusive A or T nucleotides and are not 7meG-forming sequences.

Damage formation was further analyzed in the TF core motif and its immediately adjacent DNA (20 bp on each side of the motif midpoint). This analysis shows that 7meG formation was strongly suppressed in the conserved regions of the motif sequences (Fig. 1D, 1E and Supplemental Fig. S2B) where Abf1 and Reb1 proteins directly contact DNA (Jaiswal et al., 2016; McBroom and Sadowski, 1994a). In contrast, 7meG damage levels were not affected outside of the core motif (e.g., −20 to −10 and 10 to 20 bp relative to the motif midpoint). 7meG levels were relatively even across the ‘low-occupancy’ Reb1 binding sites (Fig. 1F and Supplemental Fig. S2B), even though these sites have nearly identical motif sequence as the ‘high-occupancy’ binding sites. While damage formation was mainly suppressed in the core motif, we also saw increased 7meG levels (~1.5 fold) at a few positions (e.g., −7, −3, −2, and 0) at the edge of the Abf1 motif or between the two highly conserved regions within the motif (Fig. 1D). Moreover, analysis of A reads indicates that 3meA formation was increased at the −3 position of the ‘high-occupancy’ Reb1 sites, but not at the same position of the ‘low-occupancy’ Reb1 sites (Supplemental Fig. S3A and S3B). Intriguingly, the increased 3meA formation appears to be position dependent, because the adjacent −2 and −1 positions (both are conserved in A or T) did not show elevated 3meA damage formation. To understand why the −3 position is sensitive to MMS treatment, we analyzed the published Reb1-DNA complex structure (Jaiswal et al., 2016). Analysis of the structural data indicates that Reb1 protein binding causes a large curvature (~56°) in DNA and significantly compresses the minor groove near the −3 position (Supplemental Fig. S3C and S3D). These structural changes caused by Reb1 protein binding may play a role in modulating 3meA formation.

### Abf1 and Reb1 binding inhibits repair of 7meG lesions

To address how TF binding affects 7meG repair in cells, we analyzed NMP-seq data generated after repair incubation (e.g., 1 and 2 h repair). Repair analysis was conducted by normalizing 7meG lesions at each time point to the initial 7meG damage (i.e., 0 h repair). This analysis considers the variable amounts of initial damage along the motif sequence, which can conceivably impact remaining damage after repair. The normalization (i.e., damage after repair / initial damage) results in fraction of remaining damage, which is inversely correlated with DNA repair activity (Mao et al., 2017, 2016).

Our analysis indicates that repair of 7meG lesions was strongly suppressed at both Abf1 and Reb1 binding sites in wild-type (WT) cells, shown by peaks of unrepaired damage at 1 h (Supplemental Fig. S4A) and 2 h (Fig. 2A and 2B) near the TF binding midpoint. The repair suppression is mediated by TF binding, not the underlying DNA sequence, because no repair inhibition was observed at ‘low-occupancy’ Reb1 binding sites (Fig. 2C). Additionally, nucleosomes around the TF binding sites play an important role in affecting 7meG repair. Fast repair was observed in the nucleosome-depleted region around the TF binding site and linker DNA between two adjacent nucleosomes (Fig. 2A and 2B). In contrast, slow repair was found near nucleosome peaks, which is consistent with previous studies showing inhibition of BER at the nucleosome dyad center (Kennedy et al., 2019; Mao et al., 2017). A closer examination of remaining damage indicates that repair was suppressed in an ~30-40 bp DNA region including the conserved core motif and its immediately adjacent DNA (Fig. 2D and 2E). Hence, TF binding inhibits BER in a broader DNA region (both core motif and adjacent DNA) relative to its impact on NMP damage formation (mainly in the core motif).

**Figure 2.**
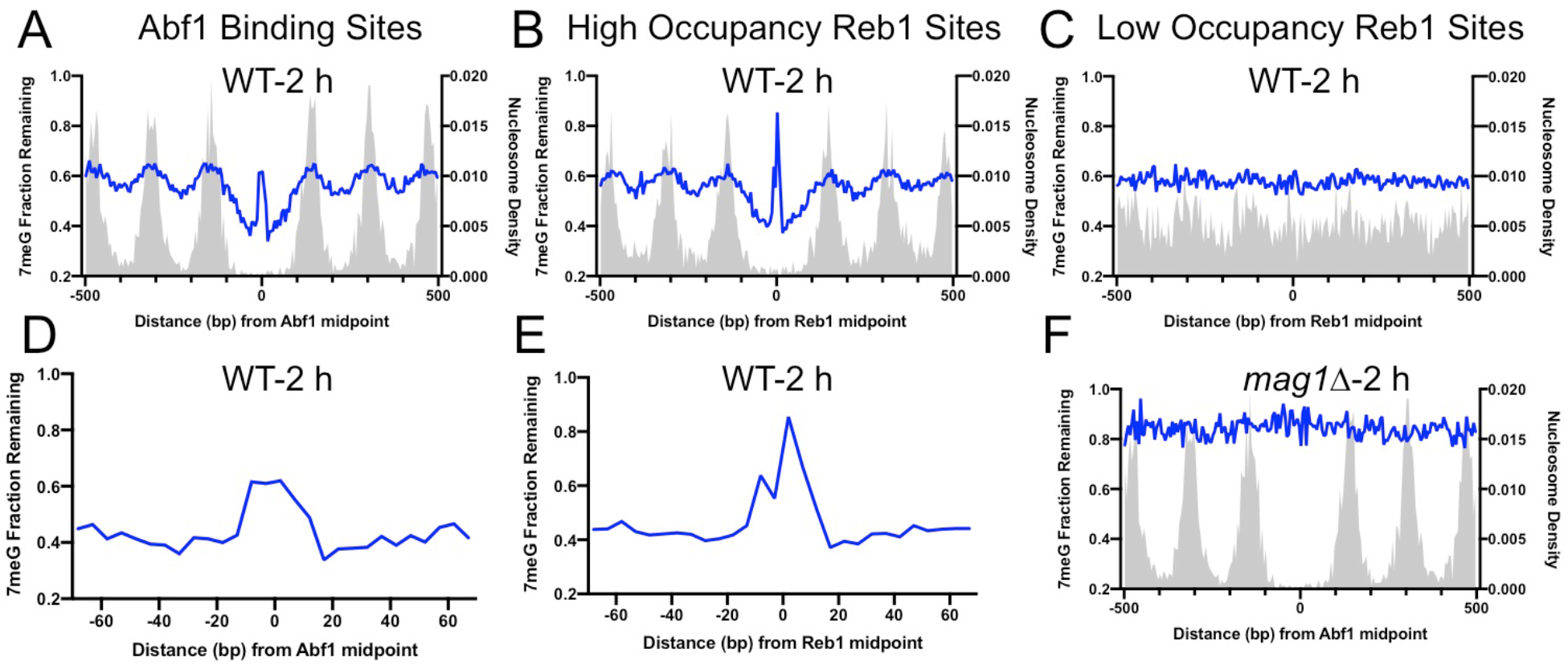
BER of 7meG lesions at Abf1 and Reb1 binding sites. **(A)** The fraction of remaining 7meG lesions (blue line) after 2 h repair at Abf1 binding sites in wild-type (WT) cells. Remaining 7meG at the binding sites and in the flanking DNA (up to 500 bp in each direction) was shown. The binding sites were obtained from the published ORGANIC method (Kasinathan et al., 2014). The plot shows the average remaining damage in 5 bp non-overlapping moving windows. The nucleosome density, which was analyzed using the published yeast MNase-seq data (Weiner et al., 2015), was plotted as the gray background. **(B)** Repair of 7meG lesions at ‘high-occupancy’ Reb1 binding sites. **(C)** Repair of 7meG at ‘low-occupancy’ Reb1 binding sites. **(D)** and **(E)** Close-up of remaining 7meG at Abf1 and ‘high-occupancy’ Reb1 sites, respectively. 7meG fraction remaining between −70 and 70 bp relative to the TF motif midpoint was shown. **(F)** The fraction of remaining 7meG lesions after 2 h repair in the *mag1*Δ mutant at Abf1 binding sites.

Repair of 7meG by BER is initiated by the Mag1 glycosylase in yeast (Wyatt et al., 1999). To test if the inhibited repair of 7meG at TF binding sites is due to reduced BER, we analyzed 7meG repair in the *mag1*Δ mutant strain. NMP-seq analysis in this mutant revealed higher levels of unrepaired 7meG lesions at 2 h than in WT (Fig. 2F), consistent with deficient BER for NMPs in the mutant. Moreover, there was no difference in remaining damage between the TF binding sites and flanking DNA in *mag1*Δ cells (Fig. 2F), confirming that BER is inhibited by TF binding.

The above analyses were performed using TF binding data generated with occupied regions of genomes from affinity-purified naturally isolated chromatin (ORGANIC), a method utilizing micrococcal nuclease (MNase) to digest native chromatin (i.e., not formaldehyde cross linked) and immunoprecipitate the TF-DNA complex for sequencing (Kasinathan et al., 2014). To confirm our findings, we used TF binding data generated with the ChIP-exonuclease (ChIP-exo) method (Rossi et al., 2021). ChIP-exo is similar to the conventional ChIP-seq, but utilizes exonuclease to cleave free DNA after chromatin immunoprecipitation to improve mapping resolution (Rhee and Pugh, 2012). Analysis of NMP-seq data at Abf1 and Reb1 ChIP-exo peaks and flanking regions showed strongly inhibited BER after 2 h repair (Supplemental Fig. S4B, left and middle panels), consistent with our analyses using ORGANIC binding data. Moreover, ChIP-exo was used to map binding sites for other yeast TFs such as Repressor Activator Protein (Rap1) (Rossi et al., 2021), an essential yeast TF involved in both activation and suppression of RNA Pol II transcription (Shore and Nasmyth, 1987). We analyzed 7meG repair at Rap1 ChIP-exo sites and found that BER was also strongly inhibited by Rap1 binding (Supplemental Fig. S4B, right panel). Hence, NMP-seq analysis using both ORGANIC and ChIP-exo binding data consistently indicates an inhibitory role of TF binding in BER.

### Depletion of Abf1 or Reb1 protein restores 7meG formation and elevates BER at their binding sites

Our data suggests that TF binding acts as a barrier to the damaging chemical MMS and BER enzymes. We hypothesize that removal of the TF would expose the binding sites to MMS and repair enzymes. As both Abf1 and Reb1 are essential for yeast survival and cannot be knocked out, we used the published Anchor-Away strategy (Haruki et al., 2008) to conditionally and rapidly export the protein from the nucleus to the cytoplasm. We then performed NMP-seq experiments in the TF-depleted yeast strains to analyze 7meG formation and repair. Both Abf1 and Reb1 anchor-away strains (Abf1-AA and Reb1-AA) were generated and used to study their impacts on gene transcription (Kubik et al., 2018, 2015). We followed the published protocol to deplete Abf1 or Reb1 from the nucleus with rapamycin. Moreover, growth of Abf1-AA or Reb1-AA strain was inhibited on rapamycin-containing plates (Supplemental Fig. S5A), confirming that nuclear depletion of either protein is lethal for yeast cells (Kubik et al., 2015).

In the control strain (WT-AA), in which no target protein is tagged for depletion, analysis of the NMP-seq data indicates that 7meG damage formation was still suppressed at the conserved motif sequences upon rapamycin treatment (Supplemental Fig. S5B), indicating that rapamycin itself had little effect on NMP damage formation. However, TF depletion in Abf1-AA or Reb1-AA cells restored damage formation at their corresponding binding sites (Supplemental Fig. S5C and S5D). For example, Abf1 depletion increased 7meG formation at Abf1 binding sites to a level comparable to the flanking DNA; however, no damage restoration was seen at Reb1 binding sites in Abf1-AA cells (Supplemental Fig. S5C). Similarly, damage was restored at Reb1 binding sites in Reb1-AA cells, but not at Abf1 sites (Supplemental Fig. S5D). Therefore, these data indicate that nuclear depletion of each TF specifically affects damage formation at its own binding sites, but has no effect on the binding sites of the other TF.

Analysis of 7meG repair in the anchor-away strains indicates that BER was restored and even elevated by removing each TF from the binding sites. Compared to the control WT-AA strain (Fig. 3A and 3B), no repair inhibition was seen at Abf1 binding sites when Abf1 was depleted (Fig. 3C). Instead, BER was faster at Abf1 binding sites relative to the flanking DNA in Abf1-AA cells (Fig. 3C), likely because these binding sites are located in nucleosome-depleted regions and damage is efficiently repaired by BER (Mao et al., 2017). Repair in the surrounding nucleosomes was also affected by Abf1 depletion (compare Fig. 3A and 3C), likely due to the weakened nucleosome organization around Abf1 binding sites in Abf1-AA cells (Kubik et al., 2018). Repair of 7meG damage was still inhibited at Reb1 binding sites in Abf1-AA cells (Fig. 3D), consistent with the notion that Reb1 protein still binds to its target sites in Abf1-AA cells. Similar results were also observed in Reb1-AA cells, where BER was inhibited at Abf1 binding sites (Fig 3E) but accelerated at Reb1 binding sites relative to the flanking DNA (Fig. 3F). Taken together, these data demonstrate that removal of Abf1 and Reb1 exposes their target sites to the damaging chemical and BER enzymes.

**Figure 3.**
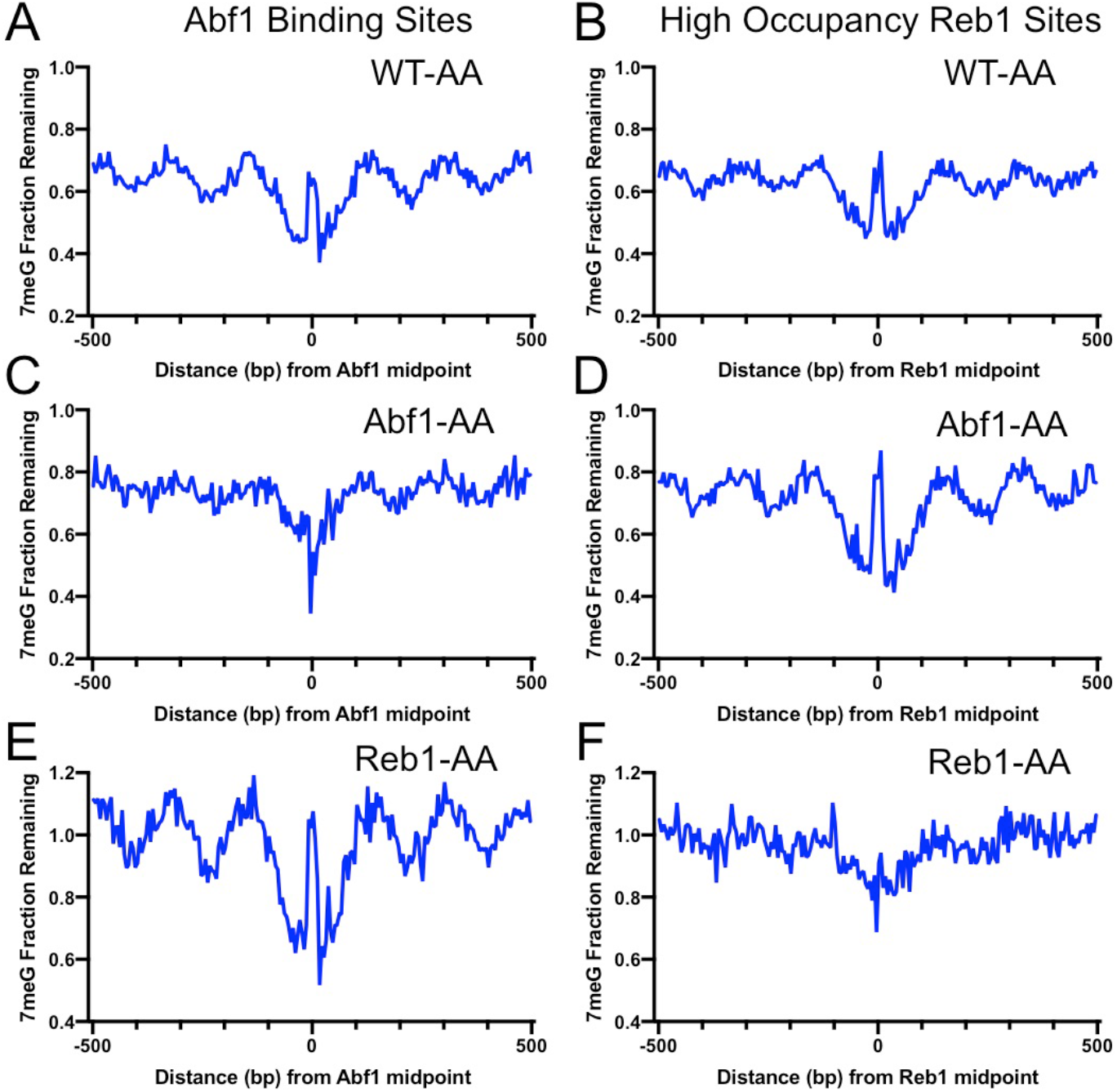
BER of 7meG lesions in anchor-away yeast strains. **(A)** Fraction of remaining 7meG lesions after 2 h repair (normalized to 0h) in rapamycin-treated WT-AA cells at Abf1 binding sites. **(B)** Remaining 7meG at ‘high-occupancy’ Reb1 binding sites in rapamycin-treated WT-AA cells. **(C)** and **(D)** Fraction of remaining 7meG at 2 h in Abf1-AA cells after rapamycin treatment at Abf1 and ‘high-occupancy’ Reb1 sites, respectively. **(E)** and **(F)** Remaining 7meG at 2 h in Reb1-AA cells after rapamycin-mediated protein depletion at Abf1 and Reb1 sites.

### Abf1 and Reb1 inhibit BER in promoters of target genes

Abf1 and Reb1 bind to the nucleosome-depleted region (NDR) of gene promoters to facilitate transcription (Kubik et al., 2018). We next sought to understand how the two TFs affect BER in the context of gene transcription. We first examined the global BER pattern by analyzing 7meG repair in WT cells for all yeast genes. Genes (n=5,205) were aligned by their transcription start site (TSS) (Park et al., 2014) and repair was analyzed in accordance with the transcriptional direction. As shown in Fig. 4A, BER (average of all genes) was generally faster in NDR relative to the coding region where DNA is organized into +1, +2, and so on nucleosomes (Fig. 4A), a pattern consistent with our previous studies (Mao et al., 2017). Hence, the global BER pattern revealed by our analysis indicates that Abf1 and Reb1 do not inhibit repair in NDR when all genes were included.

**Figure 4.**
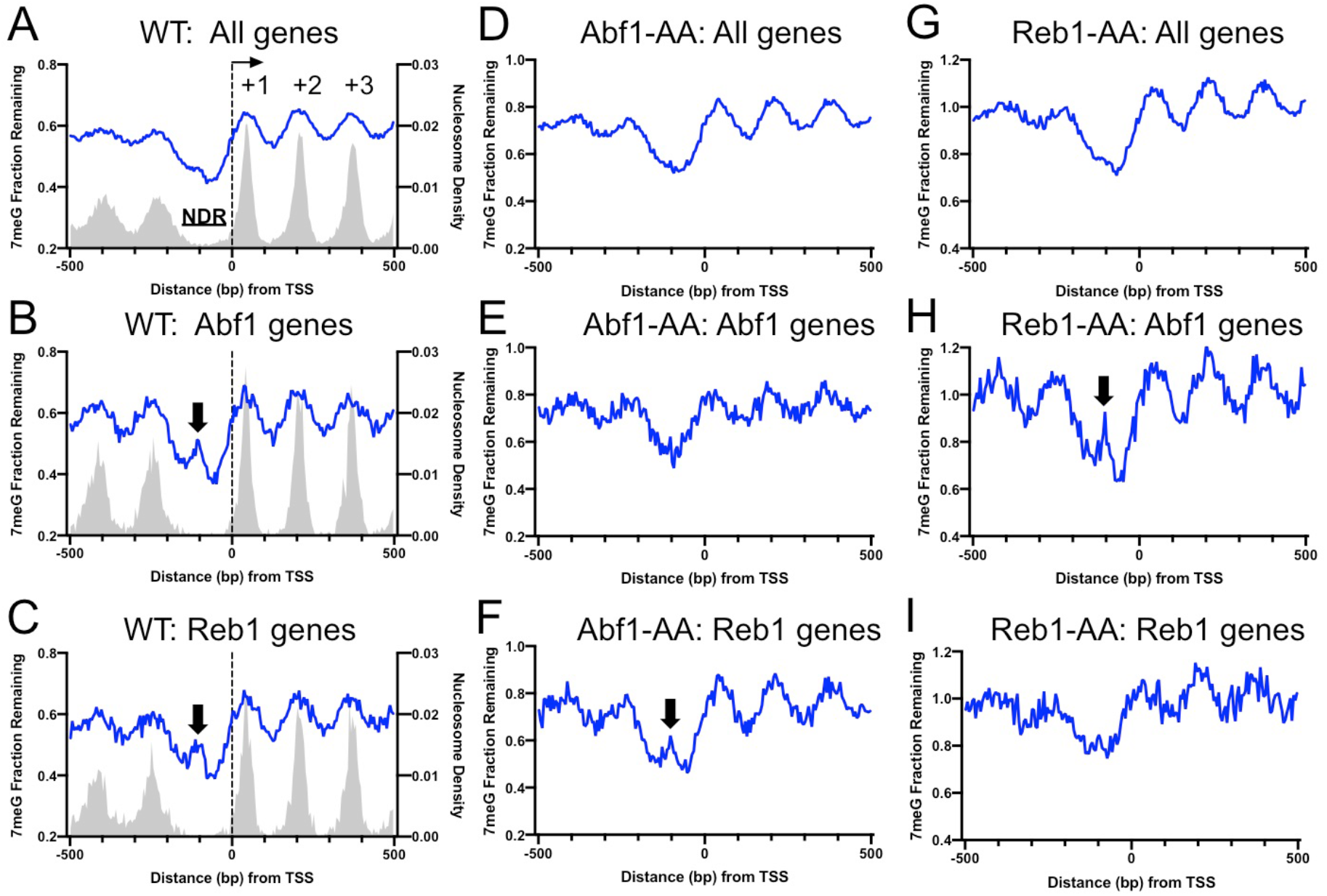
Repair of 7meG in the Abf1 and Reb1 target genes. **(A)** Average fraction of remaining 7meG lesions (blue line) after 2 h repair in all yeast genes in WT cells. Genes (n=5,205) were aligned at the TSS (position 0) and repair was plotted in accordance with gene transcriptional direction. The average damage in 5 bp moving windows is shown from upstream 500 bp to downstream 500 bp relative to the TSS. The gray background indicates nucleosome peak density. **(B)** Average fraction of remaining 7meG lesions after 2 h repair in WT cells in Abf1-linked genes (n=697). **(C)** Average fraction of remaining 7meG lesions after 2 h repair in WT cells in Reb1-linked genes (n=708). **(D)** to **(F)** Fraction of remaining 7meG at 2 h in Abf1-depleted cells for all genes, Abf1-linked, and Reb1-linked genes. **(G)** to **(I)** Fraction of remaining 7meG at 2 h in Reb1-depleted cells for all genes, Abf1-linked, and Reb1-linked genes.

As Abf1 or Reb1 do not affect BER globally, we hypothesized that they may specifically affect BER in target genes. To test this hypothesis, we linked Abf1 and Reb1 binding sites to the closest TSS of annotated genes (Park et al., 2014). This association identified 697 Abf1-linked and 708 Reb1-linked genes (see Methods for detail). We then aligned Abf1-linked and Reb1-linked genes at their TSS and plotted 7meG repair in accordance with the transcriptional direction. For each subset of genes (i.e., Abf1-linked or Reb1-linked genes), we found a prominent damage peak in NDR after 2 h repair in WT cells (Fig 4B and 4C, black arrows). The damage peak was located ~100 bp upstream of the TSS and overlapped with Abf1 or Reb1 binding peak (Supplemental Fig. S6A and S6B), suggesting that Abf1 and Reb1 indeed inhibit BER in their target promoters. This finding was further confirmed by analyzing NMP-seq data generated in the anchor-away cells. We found that depletion of Abf1 in Abf1-AA cells did not change the global BER pattern when all genes were included (Fig. 4D), but it restored repair in the NDR of Abf1 target genes (Fig. 4E). As expected, repair in Reb1 target genes was still inhibited in the Abf1-AA cells (Fig. 4F, black arrow). Similar results were found in the Reb1-AA cells (Fig. 4G to 4I). The damage peaks in NDR were not as high as repair analysis at the mapped TF binding sites (e.g., compare Fig. 4B with Fig. 2A), likely because the gene analysis was performed in each subset of genes aligned on their TSS, not the midpoint of the TF binding sites. In summary, these data indicate that Abf1 and Reb1 inhibit BER in their target promoters.

### Repair of 3meA is inhibited by TF binding in vivo and in vitro

Although 3meA is much less abundant than 7meG in MMS-treated cells, 3meA has long been known to be cytotoxic (Fu et al., 2012; Plosky et al., 2008). Conventional methods studying cellular repair of MMS-induced damage (e.g., AAG/APE1 digestion followed by gel electrophoresis) (Czaja et al., 2014) cannot distinguish repair of 7meG and 3meA. Additionally, 3meA is unstable and difficult to be synthesized *in vitro*. As NMP-seq maps both 3meA and 7meG lesions, we extracted A reads to specifically analyze 3meA repair.

Analysis of 3meA lesions in WT cells indicates that the repair was inhibited near the center of Abf1 and ‘high-occupancy’ Reb1 binding sites, as shown by high levels of remaining 3meA lesions at 2 h (Fig. 5A and 5B). In contrast, 3meA repair was not inhibited at ‘low-occupancy’ Reb1 binding sites (Fig. 5C). Interestingly, the 3meA peaks appear to be narrower than the 7meG peaks, and no clear 3meA repair inhibition was seen in nucleosomes surrounding the TF binding sites. These differences are consistent with the greater activity of Mag1 and its homologs in removing 3meA than 7meG (Connor et al., 2005), which may lead to less repair inhibition to 3meA lesions by DNA-binding proteins.

**Figure 5.**
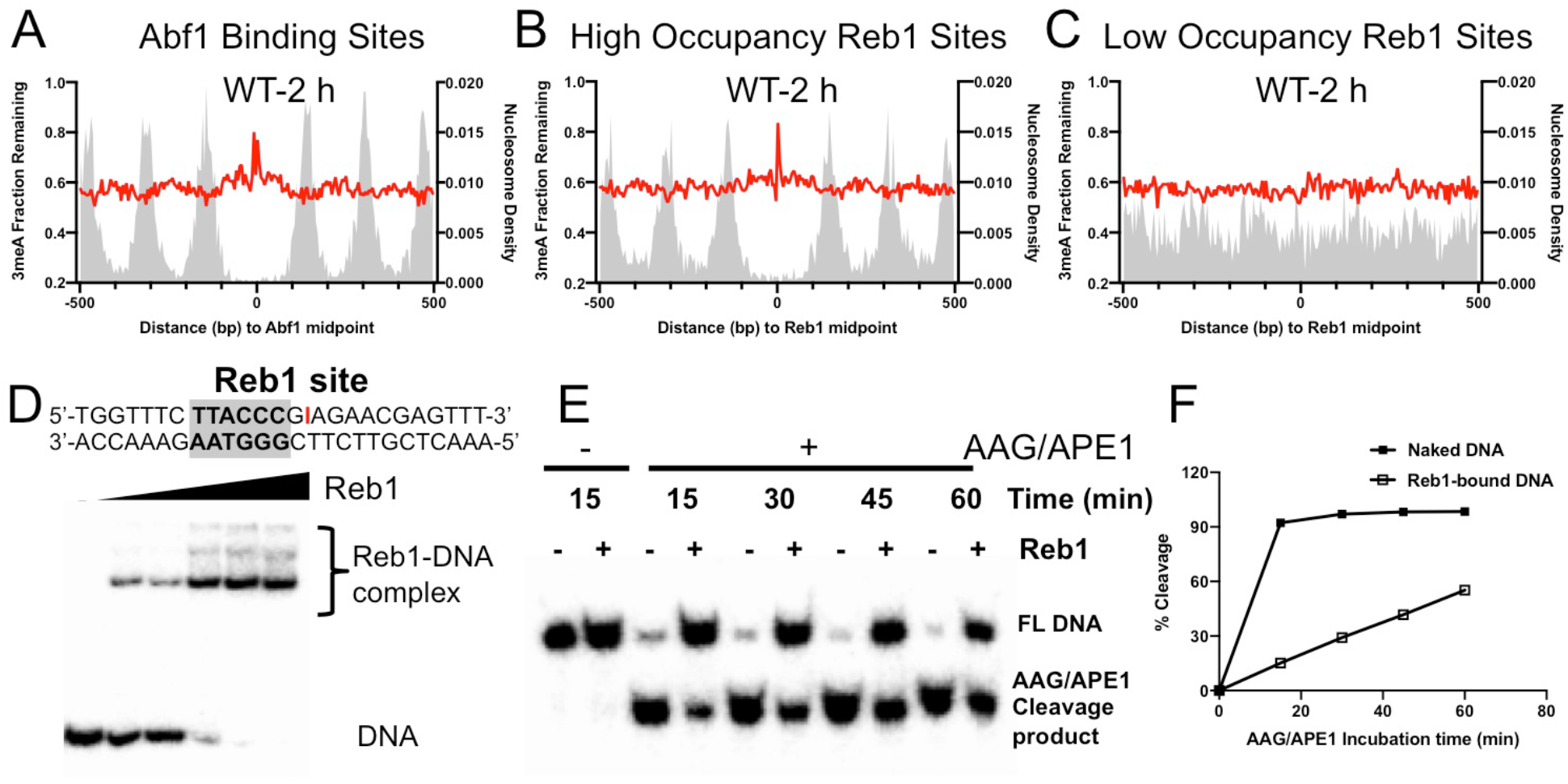
Repair of 3meA at TF binding sites. **(A)** Average fraction of remaining 3meA lesions (red line) at Abf1 binding sites mapped with the ORGANIC method. Data shows fraction of remaining 3meA lesions in 5 bp non-overlapping moving windows along the binding sites in WT cells at 2 h. **(B)** and **(C)** Fraction of remaining 3meA damage at ‘high-occupancy’ and ‘low-occupancy’ Reb1 binding sites, respectively. **(D)** The upper panel shows synthesized double-stranded DNA containing a Reb1 binding site. The inosine damage (red) was incorporated at the +4 position on the Reb1 motif strand. The lower panel shows gel shift data with DNA alone or DNA incubated with increasing amounts of purified Reb1 protein. DNA was labeled with ^32^P on the 5’ end of the motif strand. **(E)** Cleavage of the inosine-containing DNA or DNA complexed with Reb1 protein by AAG/APE1 enzymes. The substrates (naked DNA or DNA-Reb1 complex) were incubated with AAG and APE1 enzymes to cleave the damage site. DNA was analyzed on denaturing polyacrylamide gels to separate the full-length DNA (FL DNA) and the cleavage product. **(F)** Quantification of the repair gel. Graph shows the percent of cleaved DNA (lower band) relative to total DNA (lower and upper bands) at different incubation time points.

A closer examination of 3meA repair at ‘high-occupancy’ Reb1 binding sites revealed a slow repair spot at the +4 position (Supplemental Fig. S7A). Repair of 7meG was also inhibited at the same location (Supplemental Fig. S7B), suggesting that the +4 position is refractory to BER enzymes. Although the sequence at +4 position is not conserved in the Reb1 motif, the Reb1-DNA crystal structure (Jaiswal et al., 2016) shows that this position is directly contacted by the DNA binding domain of Reb1 protein (Supplemental Fig. S3C). The strong repair inhibition at the +4 position led us to further investigate BER using an *in vitro* system. To simulate 3meA repair at the Reb1 binding site, we incorporated a stable 3meA analog, inosine (denoted as I), at the +4 position of the motif strand (Fig. 5D). Inosine can naturally arise from adenine deamination in cells and is repaired by Mag1-mediated BER (Alseth et al., 2014). We found that inosine incorporation did not change Reb1 binding affinity compared to DNA without inosine (Supplemental Fig. S7C). AAG and APE1 enzymes were added to naked DNA or DNA pre-bound with purified Reb1 protein to examine BER activity *in vitro*. The AAG/APE1 cleavage product (i.e., the lower band) was analyzed in a time-course experiment to compare BER activity between free DNA and Reb1-bound DNA (Fig 5E). Quantification of the gel showed significantly reduced repair activity at the binding site in Reb1-bound DNA relative to the naked DNA substrate (Fig 5F). Reduced BER activity was also observed when inosine was placed on the other strand at the +4 position (Supplemental Fig. S7D and S7E). Hence, these *in vitro* data, consistent with our cellular damage sequencing data, indicate that BER of 3meA is suppressed by TF binding.

### BER inhibition at TF binding sites is different from NER inhibition

TF binding has been shown to inhibit NER of UV damage (Frigola et al., 2021; Hu et al., 2017); however, it is not known if NER and BER are inhibited to the same extent. Using a UV damage mapping method cyclobutane pyrimidine dimer sequencing (CPD-seq), we previously showed that formation of UV-induced CPDs is significantly suppressed at Abf1 and Reb1 binding sites (Mao et al., 2016). To investigate NER at Abf1 and Reb1 binding sites, we analyzed CPD-seq data generated in UV-irradiated yeast cells. We found that repair of CPDs at 2 h (normalized to CPDs at 0 h) was inhibited at both Abf1 and Reb1 binding sites in WT cells, shown by high levels of unrepaired CPDs at the binding sites relative to the flanking nucleosome-occupied DNA (Fig. 6A and 6B). As both Abf1 and Reb1 binding sites are localized in gene promoters (Supplemental Fig. S6), transcription-coupled NER (TC-NER) may play a role in the removal of CPDs in transcribed regions surrounding the binding sites. To reduce the interference from TC-NER, we analyzed CPD-seq data generated in a *rad26*Δ mutant strain in which TC-NER is severely diminished (Duan et al., 2020), thus allowing us to focus on global genomic NER (GG-NER). Our data indicates that GG-NER was suppressed at the center of Abf1 and Reb1 binding sites, but elevated in DNA adjacent to the center due to depletion of nucleosomes (Fig. 6C and 6D), similar to the BER pattern (Fig. 2A and 2B). Additionally, GG-NER was also modulated by nucleosomes positioned around the TF binding sites. These analyses indicate that GG-NER is inhibited by both Abf1 and Reb1 at their binding sites.

**Figure 6.**
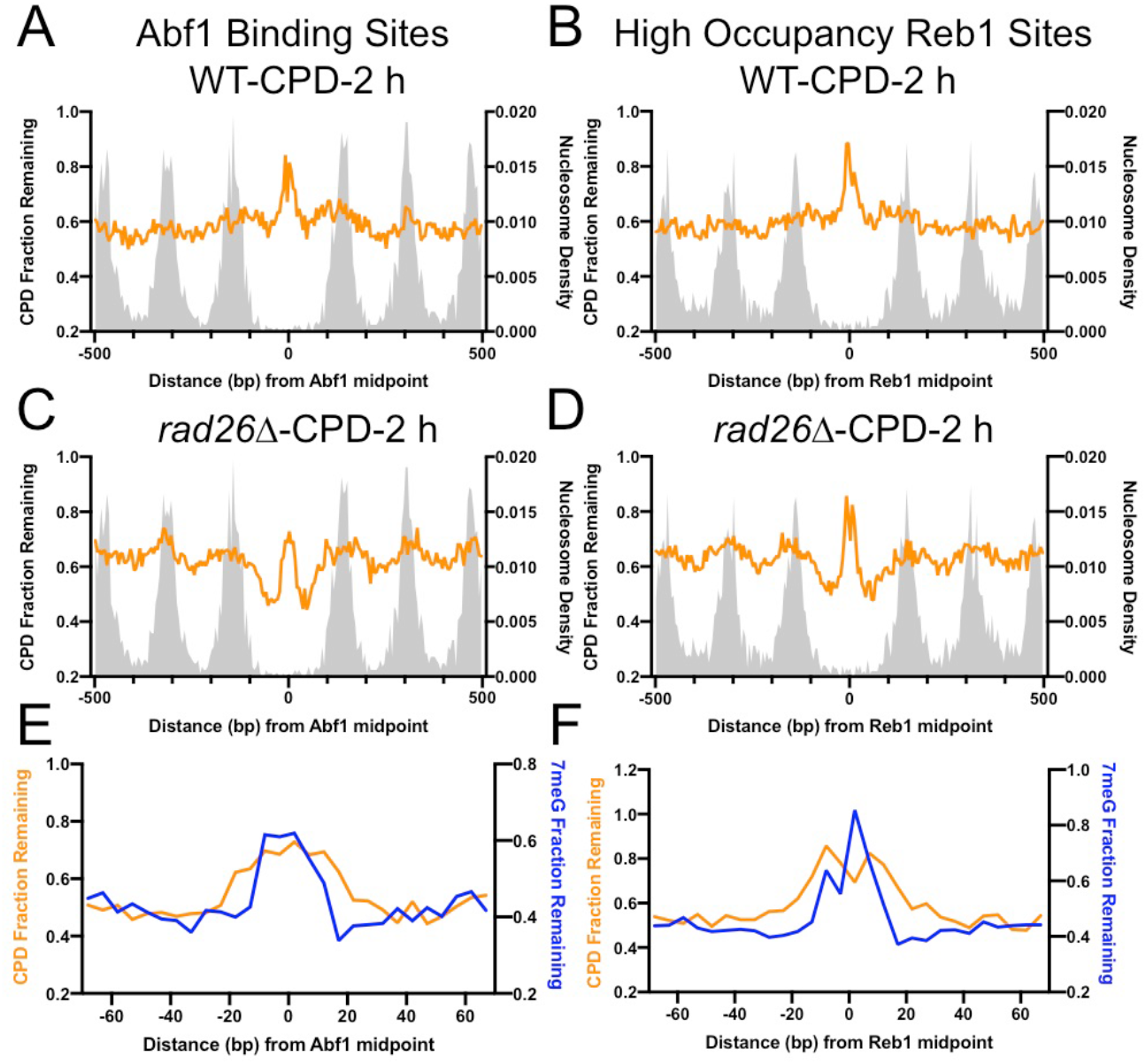
Comparison of CPD and 7meG repair at TF binding sites. **(A)** Fraction of remaining CPDs at Abf1 binding sites in WT-2 h cells. Similar to NMP-seq data analysis, the number of CPD-seq reads at 2 h was normalized to initial damage reads at 0 h. The resulting fraction of remaining CPDs was plotted at Abf1 binding sites and flanking DNA up to 500 bp. The average remaining damage in 5 bp non-overlapping moving windows was shown. **(B)** Fraction of remaining CPDs was analyzed at ‘high-occupancy’ Reb1 binding sites. **(C)** Fraction of remaining CPDs at Abf1 binding sites in the *rad26*Δ mutant strain, in which CPD repair is mainly conducted by GG-NER. **(D)** Fraction of remaining CPDs at Reb1 binding sites in the *rad26*Δ mutant cells. **(E)** Comparison between GG-NER (orange line) and BER (blue line) at Abf1 binding sites. GG-NER was analyzed using CPD-seq data (2 h relative to 0 h) generated in *rad26*Δ cells. BER analysis was conducted with NMP-seq data.

As the GG-NER pattern at the TF binding sites resembles the BER pattern revealed by our NMP-seq data, we sought to understand if the size of the inhibited DNA region is the same for both repair pathways. A comparison between CPD and 7meG repair indicates that GG-NER was inhibited in a broader DNA region at Abf1 and Reb1 binding (Fig. 6E and 6F). While BER (i.e., 7meG repair) was inhibited in ~30-40 bp DNA surrounding the center of the binding motif, inhibition of GG-NER was extended by an additional 10 bp on each side (Fig. 6E and 6F). These high-resolution sequencing data demonstrate the difference between BER and GG-NER at TF binding sites, which is consistent with the different mechanisms underling NER and BER (see Discussion).

## Discussion

In this study, we used MMS-induced damage as a model lesion and analyzed base damage distribution and BER at the binding sites of yeast TFs. Our high-resolution damage mapping data revealed an important role for TF binding in modulating initial damage formation and inhibiting BER. As base damage (e.g., oxidative, alkylation, uracil, and AP sites) has long been recognized as an important source of DNA mutations in human cancers (Maynard et al., 2009; Tubbs and Nussenzweig, 2017), the interplay between TF binding, base damage formation, and BER revealed by our study has important implications for understanding mutations in gene regulatory regions.

Our data shows that TF binding can significantly modulate NMP damage formation. Depending on the location and the conservation level in the binding motif, TF binding can both suppress and elevate damage levels. The highly conserved nucleotides in the core motif of both Abf1 and Reb1 binding sites mainly suppressed NMP damage formation (Fig. 1). NMP damage is formed via the chemical reaction between the alkylating agent and individual nucleotides (Fu et al., 2012). Due to protein-DNA interactions, nucleotides with restrained reactivity with MMS will be less sensitive and thus generate reduced amounts of damage. The highly conserved nucleotides in the Abf1 and Reb1 motifs are directly contacted by specific amino acids of the protein (Jaiswal et al., 2016; McBroom and Sadowski, 1994a). This suggests that Abf1 and Reb1 reduce the reactivity of the bound nucleotides, thus protecting conserved parts of the core motif from alkylation DNA damage. The protective role of TFs does not seem be specific for alkylation damage. UV damage formation was also reported to be suppressed by TF binding in yeast (Mao et al., 2016) and human cells (Frigola et al., 2021). Thus, TF binding may function as important mechanism in cells that protects conserved regulatory sequences from being damaged and mutated.

A few specific positions in the Abf1 motif exhibit elevated 7meG formation. Moreover, we also found a specific 3meA hotspot in the Reb1 motif. While the detailed mechanism for elevated alkylation damage formation is unclear, previous studies of UV damage formation revealed that TF binding-mediated DNA structural change plays a critical role in dictating damage yields. Indeed, human ETS (E26 transformation-specific) TFs have been shown to change the DNA geometry at their binding sites and cause individual UV damage and mutation hotspots (Elliott et al., 2018; Mao et al., 2018). Yeast Abf1 protein has been shown to bend DNA toward its minor groove (McBroom and Sadowski, 1994b). DNA bending caused by Abf1 may expose certain bases in the motif and increase their reactivity with MMS, resulting in elevated damage yields. The published complex structure of Reb1 (from *Schizosaccharomyces pombe*) with DNA (Jaiswal et al., 2016) provided an opportunity to investigate how TF-DNA interactions could modulate alkylation damage distribution. The DNA-binding domain (DBD) of Reb1 winds around two turns of duplex DNA as a series of four helix-turn-helix (HTH) domains, forming a so-called “saddle”-shaped structure (Supplemental Fig. S3C). Two homologous HTH domains, termed MybAD1 and MybAD2, are followed by two homologous repeat domains MybR1 and MybR2. C-terminal to the DBD is a transcription termination domain (TTD) that is not essential to DNA binding (Jaiswal et al., 2016). Within the central core of the Reb1 consensus (5’-GG<u>G</u>TAA-3’; the underlined G is position 0), positions +2 to 0 (i.e., GGG) are directly bound by both MybAD2 and MybR1 and exhibit significantly reduced 7meG formation (Fig. 1E). Positions −1 to −3 (i.e., TAA) are sandwiched between the subsites for MybAD1 and MybAD2, which insert recognition helices into the adjacent DNA major groove. As a result, the minor groove from positions −1 to −3 is strongly compressed in width and increased in depth (Supplemental Fig. S3D). These results suggest that preferential formation of 3meA at position −3 may be facilitated by enhanced minor groove narrowing and DNA curvature by Reb1 binding.

Our data further revealed strong inhibition of BER at Abf1 and Reb1 binding sites. Compared to NMP damage formation, repair of 7meG lesions was inhibited in a wider DNA region consisting of the core motif and some of the flanking DNA. As mentioned earlier, the conserved nucleotides in the core motif are bound by the TFs and BER enzymes could be sterically hindered to access these sites. Even the less conserved nucleotides in the core motif are also likely inaccessible to BER enzymes, since the TF protein covers the whole motif region. In addition to the core motif, structural data indicates that some nucleotides in the flanking DNA are bound by the Reb1 protein (Jaiswal et al., 2016). Although Abf1-DNA complex structure data is currently unavailable, it is conceivable that Abf1 also binds to part of the flanking DNA. The strength of protein-DNA interaction in the flanking DNA may not be as high as in the core motif, which still allows damage formation to occur, but it considerably reduces the access of BER enzymes, particularly in DNA immediately adjacent to the core motif. As BER is generally inhibited in TF-bound DNA, damage hotspots induced by TF binding cannot be efficiently repaired and may eventually cause individual mutation hotspots when DNA is replicated. Considering the conserved damage formation and repair mechanisms between yeast and human cells, our findings provide a potential explanation to mutation hotspots at TF binding sites in non-UV exposed tumors.

The comparison between NMP and CPD repair at TF binding sites provides new insights into how TFs affect BER and NER differently. While TF binding inhibits both BER and NER, we found that the affected DNA region is considerably broader in NER compared to BER. NER is inhibited in about 50-60 bp DNA centered on the midpoint of Abf1 or Reb1 binding sites, whereas BER is suppressed in a narrower DNA region (Fig. 6). The extended inhibition region in NER is consistent with more proteins being involved in NER compared to BER. Moreover, NER requires repair endonucleases to cleave upstream of the 5’ side and downstream of the 3’ side relative to the lesion, releasing a repair intermediate of ~25 nt (Huang et al., 1992; Schärer, 2013). Although UV damage located outside of the TF binding site may be recognizable by the damage recognition factor such as XPC or yeast Rad4, one of the two repair cleavage sites may still be located within the binding motif and is inaccessible to the repair endonuclease. Hence, the unique ‘dual-incision’ mechanism of NER is consistent with the broader repair-resistant DNA region around a TF binding site compared to BER.

In summary, we generated high-resolution alkylation damage and BER maps at yeast TF binding sites, which allows us to elucidate how TF binding modulates base damage formation and repair. Considering the potential connection between base damage, BER, and mutations in non-UV exposed tumors, these analyses provide important insights into cancer mutations frequently elevated at TF binding sites.

## Materials and Methods

### Yeast strains

Wild-type (WT) and *mag1*Δ strains were in the BY4741 background. The anchor-away (AA) strains, including WT-AA, Abf1-AA, and Reb1-AA, were gifts from Dr. David Shore (Kubik et al., 2018, 2015).

### MMS treatment

Yeast cells were grown in YPD (yeast extract-peptone-dextrose) medium to mid-log phase and treated with 0.4% (v/v) MMS (Acros Organics, AC15689) for 10 min to induce alkylation damage. Cells were centrifuged and washed with sterile deionized water to remove MMS. Cells were resuspended in pre-warmed YPD medium and incubated for repair in a 30 °C shaker.

The anchor-away yeast cells were pre-treated with 1 μg/ml rapamycin (Fisher Scientific, NC0678468) for 1h in YPD medium, as described in previous studies(Haruki et al., 2008; Kubik et al., 2018). At the end of rapamycin treatment, MMS was added to the culture and incubated for 10 min. After MMS treatment, cells were spun down and washed with sterile water to remove MMS. Cells were then resuspended in fresh YPD containing 1 μg/ml rapamycin for repair time points.

To damage naked yeast DNA with MMS, genomic DNA was first isolated from WT yeast cells without MMS treatment. All proteins were removed during DNA isolation by using vigorous phenol chloroform extraction, followed by ethanol precipitation. The purified DNA was incubated with MMS for 10 min. After MMS treatment, DNA was purified by phenol chloroform extraction and ethanol precipitation.

### NMP-seq library preparation

NMP-seq library preparation was described in our previous study (Mao et al., 2017). Genomic DNA was sonicated to small fragments and ligated to the first adaptor DNA. The ligation product was purified and incubated with terminal transferase and dideoxy-ATP (ddATP) to block all free 3’ ends (Ding et al., 2015). The NMP lesion site was cleaved by hAAG (NEB, M0313S) and APE1 (NEB, M0282S) to generate a new ligatable 3’ end. DNA was denatured at 95 °C and cooled on ice, followed by ligation to the second adaptor. After purification with Streptavidin beads (Thermo Fisher, 11205D), the library DNA was briefly amplified by PCR with two primers complementary to the two adaptors. Sequencing of NMP-seq libraries was conducted on an Iron Torrent platform.

### TF binding data sets

We used published yeast TF binding data sets in this study. Most analyses were performed using the published ORGANIC binding data (Kasinathan et al., 2014). Binding sites were obtained from experiments using 10 min micrococcal nuclease digestion with 80 mM NaCl, as described in our previous study (Mao et al., 2016). Only binding sites with the canonical Abf1 or Reb1 motif sequence (CGTNNNNNRNKA and TTACCC, respectively) were used for damage and repair analysis. Binding sites that did not match the motif sequences were excluded. Reb1 binding sites were further stratified into ‘high-occupancy’ (occupancy > 10) and ‘low-occupancy’ (occupancy <=10) binding sites based on the mapped occupancy levels (Kasinathan et al., 2014).

Some of our repair analyses (e.g., Supplemental Fig. S4B) were confirmed using ChIP-exo TF binding sites. The Abf1, Reb1, and Rap1 binding peaks were determined by mapping genome-wide binding sites in TAP-tagged yeast strains (e.g., Abf1-TAP, Reb1-TAP, and Rap1-TAP) in a recent ChIP-exo study (Rossi et al., 2021). The data were downloaded from the Gene Expression Omnibus, https://www.ncbi.nlm.nih.gov/geo/ (accession number GSE147927).

To identify target genes for Abf1 and Reb1, we searched gene transcription start sites (TSS) to find the closest midpoint of Abf1 or Reb1 binding sites using the ORGANIC datasets. If the TF binding site is located within 300 bp upstream or downstream of the gene TSS, the gene is identified as a putative target gene. Some binding sites are located in the middle of two divergently transcribed genes. In this case, both genes are recognized as target genes.

### NMP-seq data analysis

Analysis of NMP-seq datasets was conducted using our published protocols (Mao et al., 2017). NMP-seq sequencing reads were demultiplxed and aligned to the yeast reference genome (sacCer3) using Bowtie 2 (Langmead and Salzberg, 2012). For each mapped read, we identified the position of its 5’ end in the genome using SAMtools (Li et al., 2009) and BEDTools (Quinlan and Hall, 2010). Based on the 5’ end position, the single nucleotide immediately upstream of the 5’ end was found and the sequence on the opposing strand was identified as the putative NMP lesion. The number of sequencing reads associated with each of the four nucleotides (e.g., A, T, C, and G) was counted to estimate the enrichment of MMS-induced NMP lesions in the sequencing libraries. G reads were typically highly enriched relative to C reads, followed by A reads.

To analyze damage formation and BER at TF binding sites, we extracted G or A reads to analyze 7meG and 3meA lesions, respectively. The number of lesions at each position around the midpoint of Abf1 or Reb1 binding sites was counted using the BEDTool intersect function. For damage formation, the cellular lesion counts were normalized to the naked DNA to account for the impact of DNA sequences on NMP lesion formation. The normalized ratio was scaled to 1.0 and plotted along the TF binding sites (e.g., Fig. 1A to 1C). Plots at single nucleotide resolution (e.g., Fig. 1D to 1E) also show scaled damage ratio between cellular and naked DNA NMP-seq data. For repair analysis, damage counts at repair time points were normalized to the initial damage at 0 h to generate fraction of remaining damage. Positions with high fraction of remaining damage are indicative of slow repair, since a large fraction of damage is not repaired at that site. Some highly conserved positions at TF binding sites do not have lesion-forming nucleotides. These positions are labeled with asterisks in single-nucleotide resolution plots (e.g., Fig. 1D). Alternatively, we analyzed the average damage in a 5-bp non-overlapping moving window to show the average damage and repair in a broader DNA region (e.g., Fig. 1A and 2A).

Some NMP-seq datasets such as mag1-0 h, WT-1 h and WT-2 h, were downloaded from our published studies (NCBI GEO, accession code GSE98031). New NMP-seq data generated in this study, including NMP data in naked DNA and in anchor-away yeast strains, have been submitted to NCBI GEO (accession code GSE183622). In some of the new samples (e.g., WT-AA, Abf1-AA-rep 2), we tried to add MMS-damaged pUC19 plasmid as spike-in control to quantify repair efficiency. Hence, the fraction of remaining damage in these samples was normalized by the pUC19 read ratio between 0 h and 2 h.

### CPD-seq datasets and analysis

Yeast CPD-seq data were downloaded from NCBI GEO (accession code GSE145911). Analysis of CPD repair at Abf1 and Reb1 binding sites was performed using the same method described in NMP-seq data analysis.

### *In vitro* Reb1 binding and BER assay

Recombinant Reb1 protein was expressed in *E.coli* cells in a pET30a(+) expression vector (a gift from Dr. David Donze at Louisiana State University). Protein was purified with Co-NTA resin and eluted using 0.25 M imidazole. The purity of the eluted protein was ~90% as judged by Coomassie-stained SDS-PAGE. The nominal molecular weight of the recombinant construct was ~55 kDa. Protein concentration was determined by UV absorption at 280 nm. Reb1-DNA binding was analyzed using electrophoretic mobility shift assay (EMSA). Inosine lesion containing oligonucleotide (40 μM), or control oligonucleotide without inosine, was labelled with γ-^32^P ATP (20 μCi) (Perkin Elmer) in a 25 μL reaction containing 1X PNK buffer and 15 units of polynucleotide kinase (New England Biolabs) by incubating at 37°C for 45 minutes. The reaction was heat inactivated at 65°C for 15 minutes. G-25 sephadex™ G-50 DNA grade resin columns were used to remove unincorporated γ-^32^P ATP according to manufacturer’s instructions (illustra™ GE Healthcare). The purified strand was used for subsequent annealing with equal amount of complementary strand in 50 μL total volume. The annealed duplex DNA (20 pmol) was mixed with increasing concentrations of Reb1 (5.5 pmol, 11 pmol, 22 pmol, 33 pmol and 44 pmol) in 50 μL reactions containing 1X EMSA buffer (see Fig. 5D). The binding reaction was incubated on ice for 40 minutes. Free DNA and DNA bound by Reb1 were loaded onto a 12% native PAGE and separated by gel electrophoresis at 200 V for 30 minutes. The gel was exposed to a phosphor screen and the image was scanned using a Typhoon FLA7000 scanner (GE Healthcare). Gel quantification was performed with the ImageQuant software (GE healthcare).

For BER assays, equal amount of naked DNA and DNA bound by Reb1 protein (~ 5pmol) were incubated with AAG (10 units) and APE1 (1 unit) (New England Biolabs) in a 20 μL reaction containing 1X Thermopol buffer (20mM Tris HCl pH 8.8, 10 mM (NH4)_2_SO_4_, 10mM KCl, 2mM MgSO_4_, 0.1% Triton X-100) at 37°C for 15, 30, 45 and 60 minutes. After BER cleavage, DNA was purified using Phenol:Chloroform:Isoamyl alcohol extraction and precipitated using ethanol. The purified DNA was resuspended in formamide (80%) and denatured at 95°C for 10 minutes. The denatured DNA was analyzed by electrophoresis at 200V for 30 min using 12% polyacrylamide urea gels. The gel was exposed to a phosphor screen and imaged using a Typhoon FLA7000 scanner and quantified by ImageQuant.

### Structural analysis

The co-crystal structure of *Schizosaccharomyces pombe* Reb1 with terminator DNA that harbors a core consensus 5’-GGGTAA-3’ (PDB: 5eyb) was used (Jaiswal et al., 2016). The bound DNA was analyzed using curves+ (Lavery et al., 2009) to fit the helical curvature and groove parameters. Values of helical parameters were reported as averages ± standard deviations for the two copies found in the asymmetric unit. Atom-centered electrostatic potentials at 25°C in implicit water were computed using APBS (Baker et al., 2001) based on atomic charges and radii assigned from the AMBER14 forcefield. The solute dielectric was set to 8 based on recently reported measurements on duplex DNA (Cuervo et al., 2014).

## Acknowledgements

We thank Mark Wildung and Wei Wei Du for technical assistance with Ion Proton sequencing. We also thank Dr. David Shore for providing anchor-away yeast strains. This work was supported by National Institute of Environmental Health Sciences Grants R21ES029302 (to P.M. and J.J.W.), R01ES028698 (to J.J.W.), NSF grant MCB 2028902 (to G. M. K. P), and a pilot grant from UNM Center for Metals in Biology and Medicine (P20GM130422). This research was partially supported by UNM Comprehensive Cancer Center Support Grant NCI P30CA118100 and UNM ATG Shared Resource.

## Supplemental Figures

**Figure S1.**
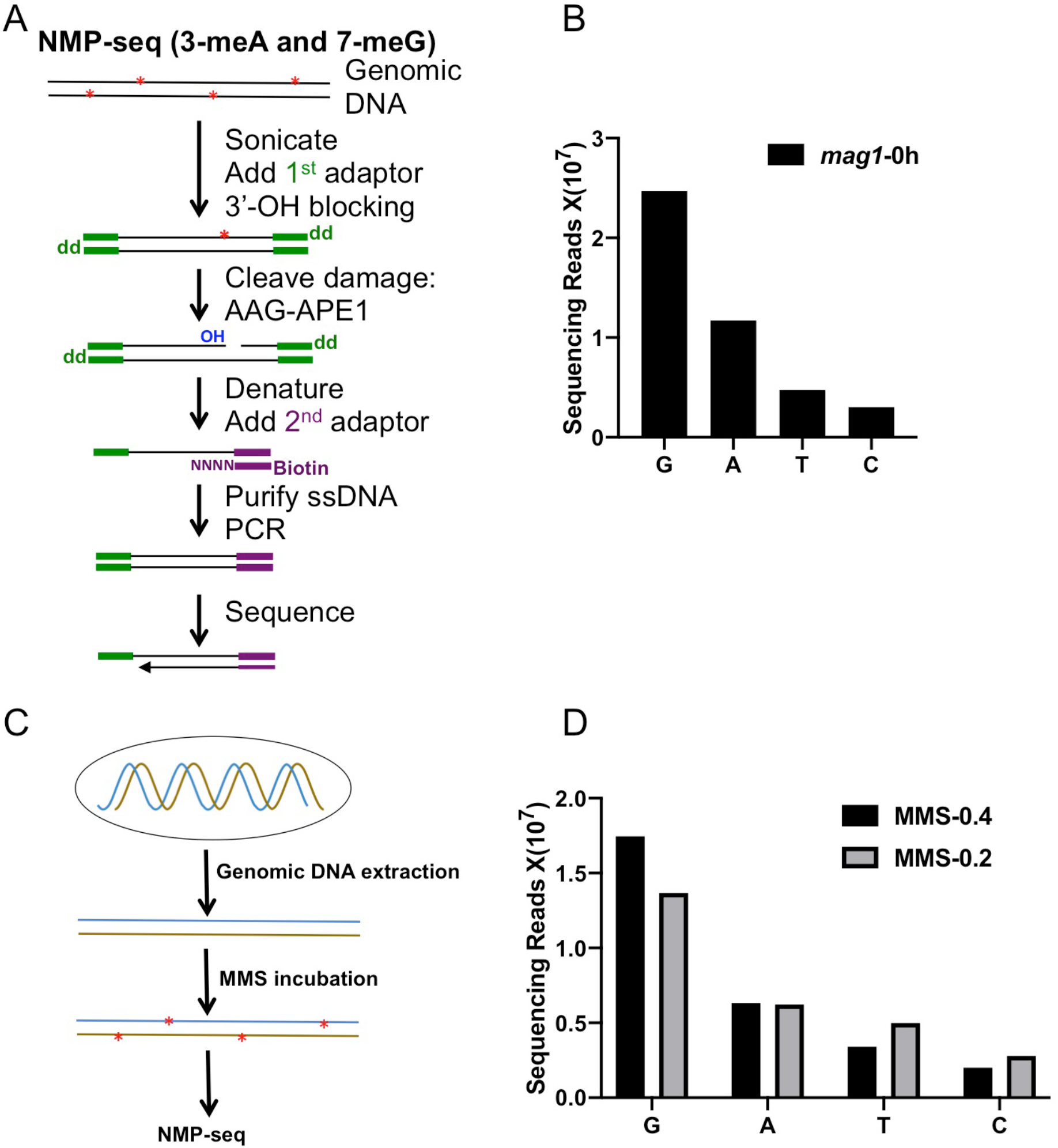
NMP-seq methodology and damage mapping in naked yeast genomic DNA. **(A)** NMP-seq methodology. Genomic DNA was first sonicated to short fragments (~400 bp) and ligated to the 1^st^ adaptor (green). After blocking all free 3’-OH groups with terminal transferase and dideoxy-ATP (dd), DNA fragments were digested with AAG and APE1 to generate a new nick with a ligatable 3’-OH group at the NMP damage site. After denaturing to obtain single-stranded DNA, the new 3’ end is ligated to a splint adaptor (2^nd^ adaptor; purple) and the ligation occurs exactly at the damage site. The biotin on the 2^nd^ adaptor allows purification of ligation product with the Streptavidin beads. The purified product is used as the template for PCR amplification, using primers complementary to the 1^st^ and 2^nd^ adaptors. The resulting library is sequenced on an Iron Torrent sequencer using a sequencing primer complementary to the 2^nd^ adaptor. **(B)** NMP-seq read counts in MMS-treated *mag1* cells (0.4% MMS for 10 min). G and A reads are associated with 7meG and 3meA lesions in the genome. **(C)** Schematic for damage mapping in naked yeast genomic DNA. It differs from mapping cellular NMP lesions by inducing damage in purified DNA, instead of cellular DNA bound by proteins. **(D)** NMP-seq read counts for naked genomic DNA treated with 0.4% or 0.2% of MMS for 10 min.

**Figure S2.**
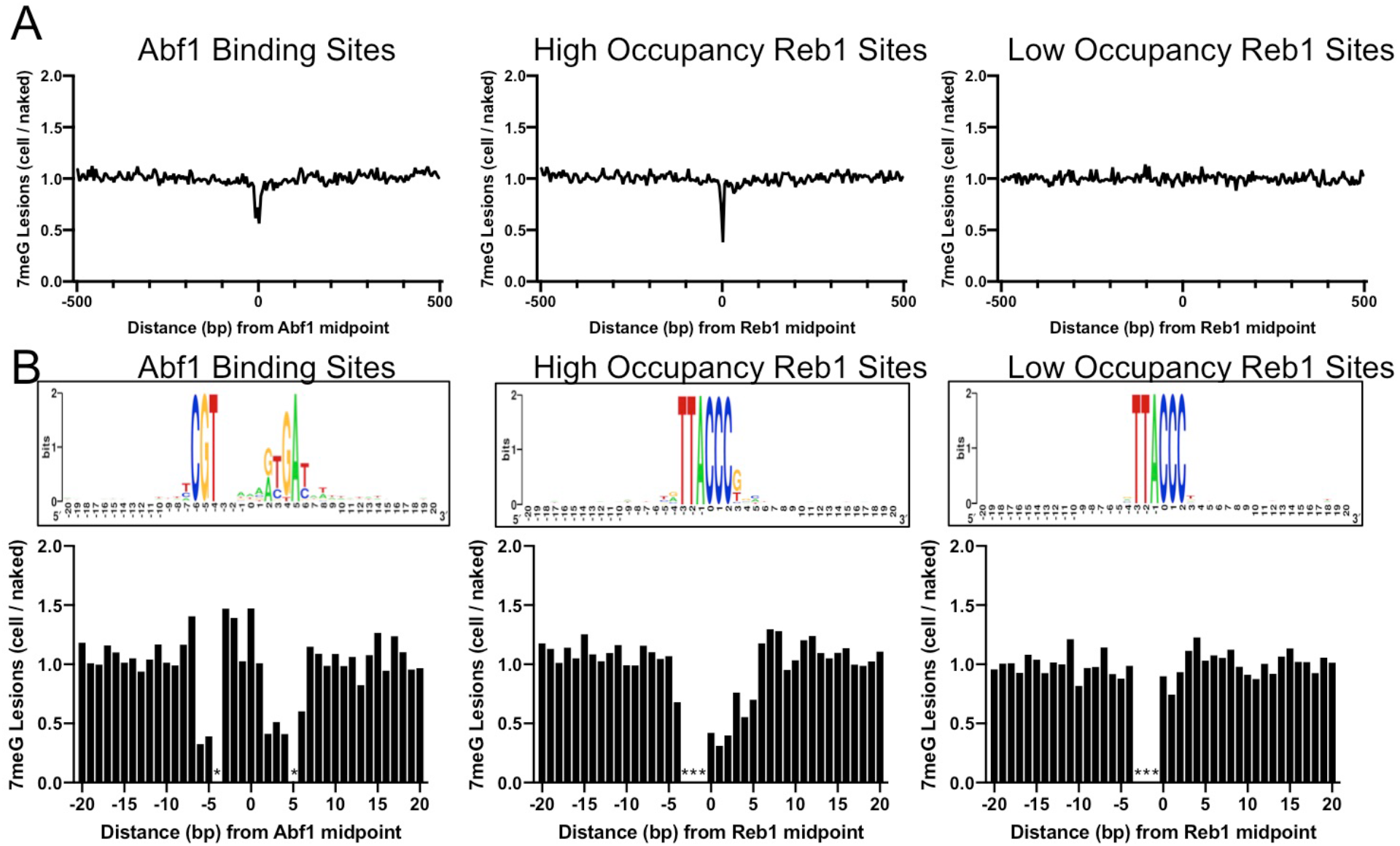
Independent repeat of 7meG formation at Abf1 and Reb1 binding sites. **(A)** The average cellular 7meG damage level (normalized to naked DNA) in a 5-bp non-overlapping moving window spanning 500 bp around the midpoint of Abf1, ‘high-occupancy’ Reb1, and ‘low-occupancy’ Reb1 sites. **(B)** Normalized 7meG damage levels at each individual position in the binding motif and its immediately adjacent DNA for Abf1, ‘high-occupancy’ Reb1, and ‘low-occupancy’ Reb1 sites.

**Figure S3.**
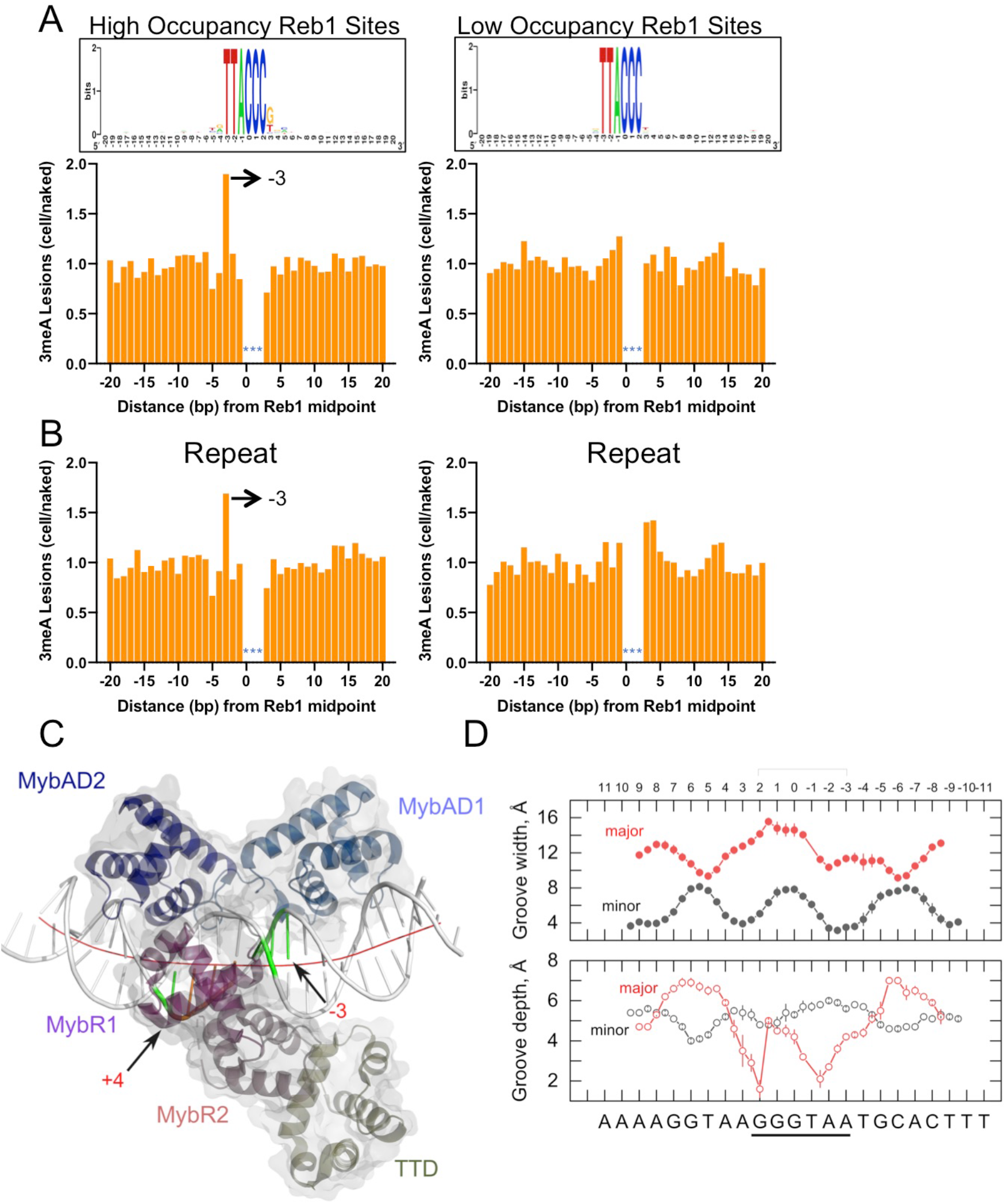
A hotspot of 3meA damage in the Reb1 motif and position-dependent distortion of Reb1-bound DNA of potential relevance to base methylation damage. **(A)** Left panel shows cellular 3meA reads (normalized to naked DNA) at ‘high-occupancy’ Reb1 binding sites. Right panel indicates 3meA reads at ‘low-occupancy’ Reb1 binding sites. Data was generated in the *mag1* mutant cells. (**B**) An independent repeat of the NMP-seq experiment in *mag1* cells showing high 3meA formation at the −3 position of Reb1 motif. Left and right panels show 3meA formation at ‘high-occupancy’ and ‘low-occupancy’ Reb1 binding sites, respectively. (**C**) One of the *S. pombe* Reb1/DNA complexes in the co-crystal structure (PDB: 5eyb). The Reb1 DNA-binding domain (DBD) was colored by domain structure, with the C-terminal transcription termination domain (TTD) also shown. Positions −3 and +4 are highlighted. The fitted curvature in helical axis is shown in red. (**D**) Widths and depths of the major and minor grooves along Reb1-bound DNA (consensus is 5’-GGGTAA-3’), reported as averages ± SD for the two complexes in the asymmetric unit.

**Figure S4.**
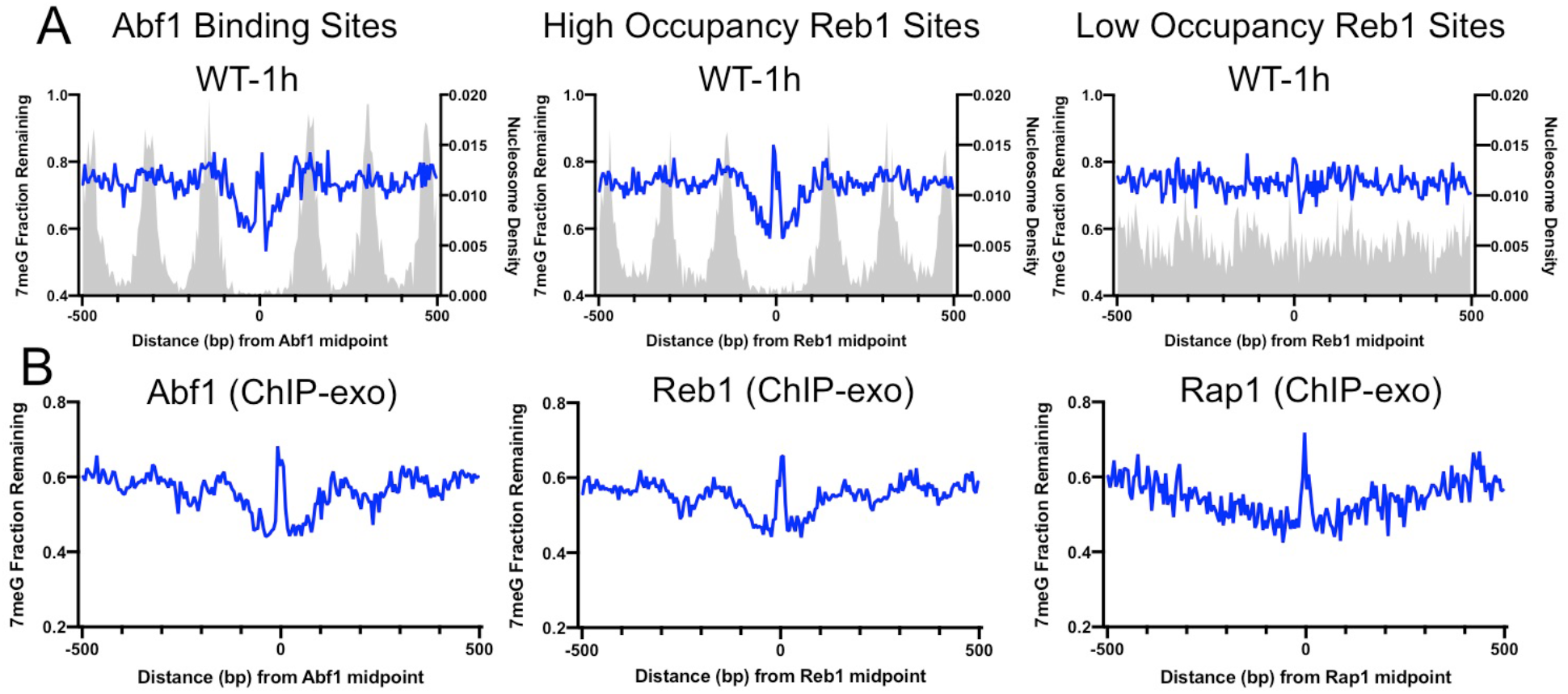
Inhibition of 7meG repair at TF binding sites mapped by ORGANIC and ChIP-exo. **(A)** Fraction of remaining 7meG lesions at 1 h in WT cells. Repair was analyzed at Abf1 (left), ‘high-occupancy’ Reb1 (middle), and ‘low-occupancy’ Reb1 (right) sites generated by ORGANIC. **(B)** Fraction of remaining 7meG lesions at 2 h in WT cells at Abf1 (left), Reb1 (middle), and Rap1 (right) binding sites identified by ChIP-exo.

**Figure S5.**
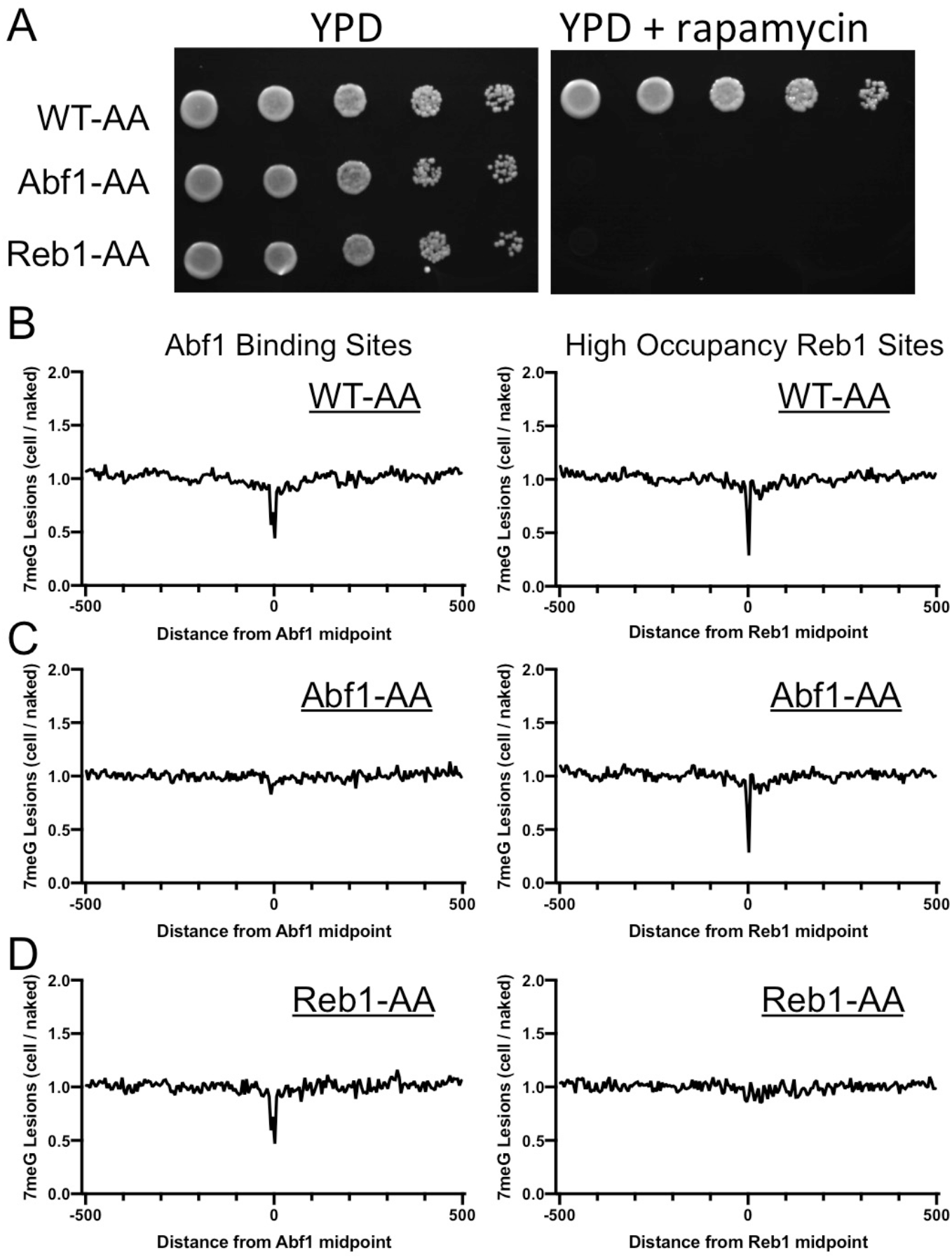
Depletion of Abf1 or Reb1 restores 7meG damage formation at the corresponding binding sites in yeast. **(A)** Lethality caused by Abf1 or Reb1 depletion on rapamycin-containing YPD plates. Yeast strains, WT-AA, Abf1-AA, and Reb1-AA, were grown on regular YPD or YPD with 1 μg/ml rapamycin. Pictures were taken after growing at 30 °C for 72 h. **(B)** to **(D)** Distribution of 7meG lesions at Abf1 (left) and Reb1 (right) binding sites in WT-AA, Abf1-AA, and Reb1-AA cells. All the three strains were pre-treated with rapamycin for 1 h, followed by MMS treatment for 10 min. Damage was mapped with NMP-seq.

**Figure S6.**
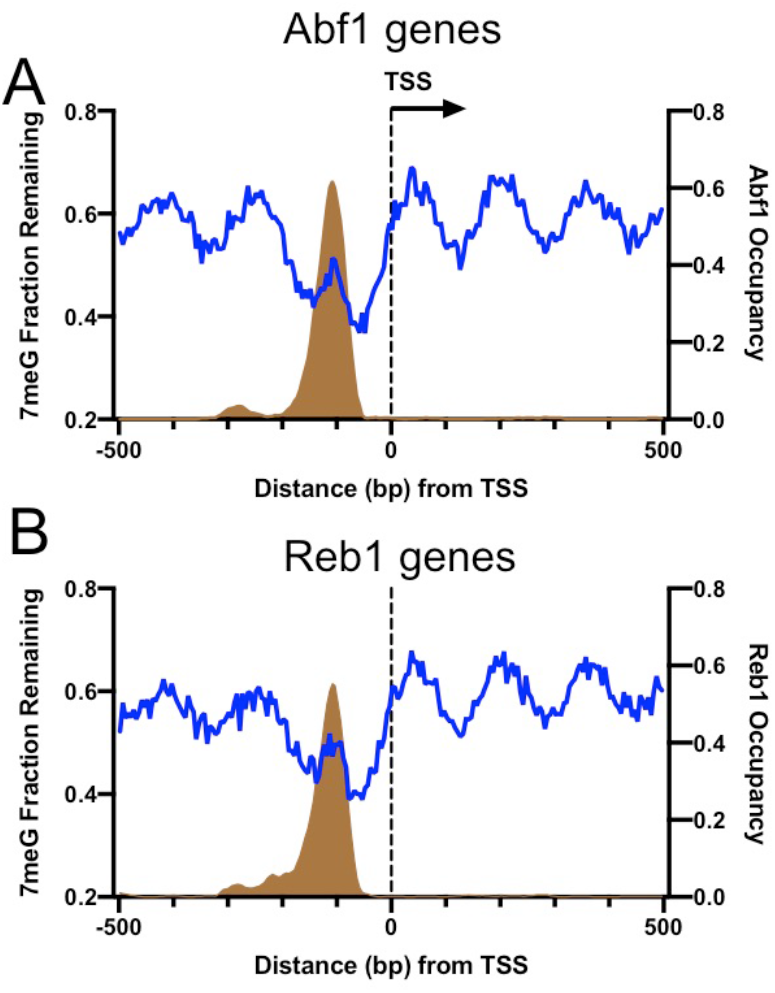
Damage peaks overlap with Abf1 or Reb1 binding sites in gene promoters. **(A)** Remaining 7meG and Abf1 occupancy in Abf1-linked genes. Blue line indicates 7meG damage and brown area depicts Abf1 occupancy. TSS stands for transcription start site. **(B)** Remaining 7meG and Reb1 occupancy in Reb1-linked genes. Genes were aligned at the TSS. Repair and TF occupancy were analyzed from 500 bp upstream to 500 bp downstream relative to the TSS.

**Figure S7.**
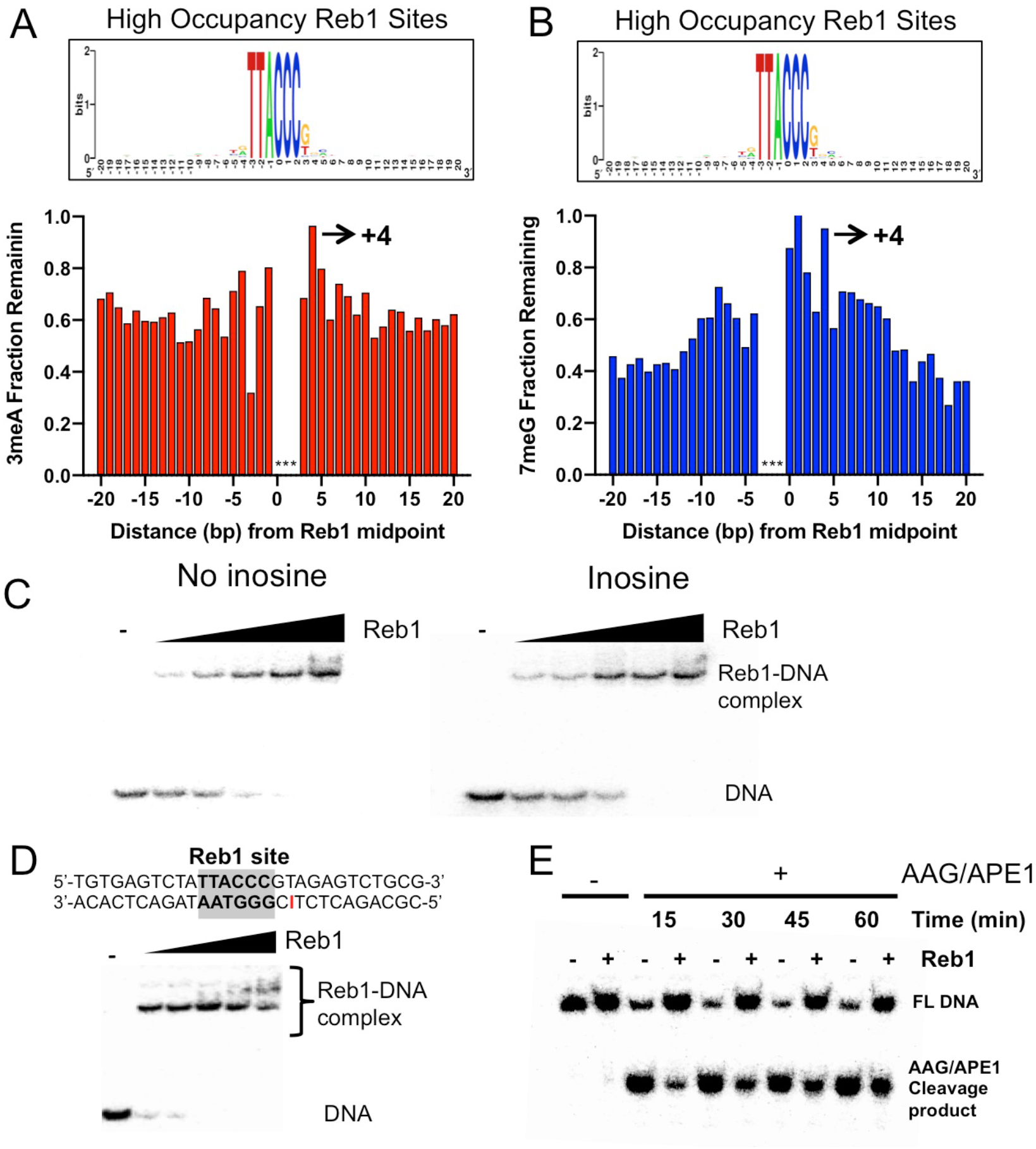
Repair inhibition at the +4 position of Reb1 binding sites. **(A)** Fraction of remaining 3meA damage at ‘high-occupancy’ Reb1 binding sites. Each bar indicates one nucleotide position in the binding motif and its adjacent DNA. The consensus motif sequence is shown on the top. **(B)** Fraction of remaining 7meG damage at ‘high-occupancy’ Reb1 binding sites. **(C)** Gel shift assays analyzing binding of synthesized double-stranded oligonucleotides with purified Reb1 protein. Left panel shows DNA without damage. Right panel shows DNA containing inosine at the +4 position. **(D)** The top panel shows DNA substrate with an inosine incorporated at the +4 position on the non-motif strand. The lower panel shows binding between the DNA and Reb1 protein. **(E)** Cleave of inosine-containing DNA (naked DNA) or DNA bound by Reb1 by AAG/APE1. The damage was placed at the +4 position of the non-motif strand.

## References

Alseth I, Dalhus B, Bjørås M. 2014. Inosine in DNA and RNA. Current Opinion in Genetics & Development, Molecular and genetic bases of disease 26:116–123. doi:10.1016/j.gde.2014.07.008

Baker NA, Sept D, Joseph S, Holst MJ, McCammon JA. 2001. Electrostatics of nanosystems: Application to microtubules and the ribosome. PNAS 98:10037–10041. doi:10.1073/pnas.181342398

Conconi A, Liu X, Koriazova L, Ackerman EJ, Smerdon MJ. 1999. Tight correlation between inhibition of DNA repair in vitro and transcription factor IIIA binding in a 5S ribosomal RNA gene. EMBO J 18:1387–1396. doi:10.1093/emboj/18.5.1387

Connor EE, Wilson JJ, Wyatt MD. 2005. Effects of Substrate Specificity on Initiating the Base Excision Repair of N-Methylpurines by Variant Human 3-Methyladenine DNA Glycosylases. Chem Res Toxicol 18:87–94. doi:10.1021/tx049822q

Cuervo A, Dans PD, Carrascosa JL, Orozco M, Gomila G, Fumagalli L. 2014. Direct measurement of the dielectric polarization properties of DNA. PNAS 111:E3624–E3630. doi:10.1073/pnas.1405702111

Czaja W, Mao P, Smerdon MJ. 2014. Chromatin remodelling complex RSC promotes base excision repair in chromatin of Saccharomyces cerevisiae. DNA Repair 16:35–43. doi:10.1016/j.dnarep.2014.01.002

Ding J, Taylor MS, Jackson AP, Reijns MAM. 2015. Genome-wide mapping of embedded ribonucleotides and other noncanonical nucleotides using emRiboSeq and EndoSeq. Nature Protocols 10:1433–1444. doi:10.1038/nprot.2015.099

Duan M, Selvam K, Wyrick JJ, Mao P. 2020. Genome-wide role of Rad26 in promoting transcription-coupled nucleotide excision repair in yeast chromatin. Proc Natl Acad Sci USA 202003868. doi:10.1073/pnas.2003868117

Elliott K, Boström M, Filges S, Lindberg M, Eynden JV den, Ståhlberg A, Clausen AR, Larsson E. 2018. Elevated pyrimidine dimer formation at distinct genomic bases underlies promoter mutation hotspots in UV-exposed cancers. PLOS Genetics 14:e1007849. doi:10.1371/journal.pgen.1007849

Friedberg EC, Walker GC, Siede W, Wood RD, Schultz RA, Ellenburger T. 2006. DNA repair and mutagenesis.DNA Repair and Mutagenesis. ASM Press.

Frigola J, Sabarinathan R, Gonzalez-Perez A, Lopez-Bigas N. 2021. Variable interplay of UV-induced DNA damage and repair at transcription factor binding sites. Nucleic Acids Research 49:891–901. doi:10.1093/nar/gkaa1219

Fu D, Calvo JA, Samson LD. 2012. Balancing repair and tolerance of DNA damage caused by alkylating agents. Nature Reviews Cancer 12:104–120. doi:10.1038/nrc3185

Guo YA, Chang MM, Huang W, Ooi WF, Xing M, Tan P, Skanderup AJ. 2018. Mutation hotspots at CTCF binding sites coupled to chromosomal instability in gastrointestinal cancers. Nature Communications 9:1520. doi:10.1038/s41467-018-03828-2

Haruki H, Nishikawa J, Laemmli UK. 2008. The Anchor-Away Technique: Rapid, Conditional Establishment of Yeast Mutant Phenotypes. Molecular Cell 31:925–932. doi:10.1016/j.molcel.2008.07.020

Hu J, Adebali O, Adar S, Sancar A. 2017. Dynamic maps of UV damage formation and repair for the human genome. PNAS 114:6758–6763. doi:10.1073/pnas.1706522114

Huang JC, Svoboda DL, Reardon JT, Sancar A. 1992. Human nucleotide excision nuclease removes thymine dimers from DNA by incising the 22nd phosphodiester bond 5’ and the 6th phosphodiester bond 3’ to the photodimer. Proc Natl Acad Sci U S A 89:3664–3668. doi:10.1073/pnas.89.8.3664

Jaiswal R, Choudhury M, Zaman S, Singh S, Santosh V, Bastia D, Escalante CR. 2016. Functional architecture of the Reb1-Ter complex of *Schizosaccharomyces pombe*. Proc Natl Acad Sci USA 113:E2267–E2276. doi:10.1073/pnas.1525465113

Jolma A, Yan J, Whitington T, Toivonen J, Nitta KR, Rastas P, Morgunova E, Enge M, Taipale M, Wei G, Palin K, Vaquerizas JM, Vincentelli R, Luscombe NM, Hughes TR, Lemaire P, Ukkonen E, Kivioja T, Taipale J. 2013. DNA-Binding Specificities of Human Transcription Factors. Cell 152:327–339. doi:10.1016/j.cell.2012.12.009

Kaiser VB, Taylor MS, Semple CA. 2016. Mutational Biases Drive Elevated Rates of Substitution at Regulatory Sites across Cancer Types. PLOS Genetics 12:e1006207. doi:10.1371/journal.pgen.1006207

Kasinathan S, Orsi GA, Zentner GE, Ahmad K, Henikoff S. 2014. High-resolution mapping of transcription factor binding sites on native chromatin. Nat Methods 11:203–209. doi:10.1038/nmeth.2766

Kennedy EE, Li C, Delaney S. 2019. Global Repair Profile of Human Alkyladenine DNA Glycosylase on Nucleosomes Reveals DNA Packaging Effects. ACS Chem Biol 14:1687–1692. doi:10.1021/acschembio.9b00263

Kondo N, Takahashi A, Ono K, Ohnishi T. 2010. DNA Damage Induced by Alkylating Agents and Repair Pathways. Journal of Nucleic Acids 2010:e543531. doi:10.4061/2010/543531

Krokan HE, Bjørås M. 2013. Base Excision Repair. Cold Spring Harb Perspect Biol 5:a012583. doi:10.1101/cshperspect.a012583

Kubik S, Bruzzone MJ, Jacquet P, Falcone J-L, Rougemont J, Shore D. 2015. Nucleosome Stability Distinguishes Two Different Promoter Types at All Protein-Coding Genes in Yeast. Molecular Cell 60:422–434. doi:10.1016/j.molcel.2015.10.002

Kubik S, O’Duibhir E, de Jonge WJ, Mattarocci S, Albert B, Falcone J-L, Bruzzone MJ, Holstege FCP, Shore D. 2018. Sequence-Directed Action of RSC Remodeler and General Regulatory Factors Modulates +1 Nucleosome Position to Facilitate Transcription. Molecular Cell 71:89–102.e5. doi:10.1016/j.molcel.2018.05.030

Langmead B, Salzberg SL. 2012. Fast gapped-read alignment with Bowtie 2. Nat Methods 9:357–359. doi:10.1038/nmeth.1923

Lavery R, Moakher M, Maddocks JH, Petkeviciute D, Zakrzewska K. 2009. Conformational analysis of nucleic acids revisited: Curves+. Nucleic Acids Res 37:5917–5929. doi:10.1093/nar/gkp608

Li H, Handsaker B, Wysoker A, Fennell T, Ruan J, Homer N, Marth G, Abecasis G, Durbin R, 1000 Genome Project Data Processing Subgroup. 2009. The Sequence Alignment/Map format and SAMtools. Bioinformatics 25:2078–2079. doi:10.1093/bioinformatics/btp352

Li M, Ko T, Li S. 2015. High-resolution Digital Mapping of N-Methylpurines in Human Cells Reveals Modulation of Their Induction and Repair by Nearest-neighbor Nucleotides. J Biol Chem 290:23148–23161. doi:10.1074/jbc.M115.676296

Mao P, Brown AJ, Esaki S, Lockwood S, Poon GMK, Smerdon MJ, Roberts SA, Wyrick JJ. 2018. ETS transcription factors induce a unique UV damage signature that drives recurrent mutagenesis in melanoma. Nature Communications 9:2626. doi:10.1038/s41467-018-05064-0

Mao P, Brown AJ, Malc EP, Mieczkowski PA, Smerdon MJ, Roberts SA, Wyrick JJ. 2017. Genome-wide maps of alkylation damage, repair, and mutagenesis in yeast reveal mechanisms of mutational heterogeneity. Genome Res 27:1674–1684. doi:10.1101/gr.225771.117

Mao P, Smerdon MJ, Roberts SA, Wyrick JJ. 2016. Chromosomal landscape of UV damage formation and repair at single-nucleotide resolution. PNAS 113:9057–9062. doi:10.1073/pnas.1606667113

Mao P, Wyrick JJ. 2019. Organization of DNA damage, excision repair, and mutagenesis in chromatin: A genomic perspective. DNA Repair, Cutting-edge Perspectives in Genomic Maintenance VI 81:102645. doi:10.1016/j.dnarep.2019.102645

Maynard S, Schurman SH, Harboe C, de Souza-Pinto NC, Bohr VA. 2009. Base excision repair of oxidative DNA damage and association with cancer and aging. Carcinogenesis 30:2–10. doi:10.1093/carcin/bgn250

McBroom LD, Sadowski PD. 1994a. Contacts of the ABF1 protein of Saccharomyces cerevisiae with a DNA binding site at MATa. J Biol Chem 269:16455–16460.

McBroom LD, Sadowski PD. 1994b. DNA bending by Saccharomyces cerevisiae ABF1 and its proteolytic fragments. Journal of Biological Chemistry 269:16461–16468. doi:10.1016/S0021-9258(17)34029-2

Melton C, Reuter JA, Spacek DV, Snyder M. 2015. Recurrent somatic mutations in regulatory regions of human cancer genomes. Nature Genetics 47:710–716. doi:10.1038/ng.3332

Morova T, McNeill DR, Lallous N, Gönen M, Dalal K, Wilson DM, Gürsoy A, Keskin Ö, Lack NA. 2020. Androgen receptor-binding sites are highly mutated in prostate cancer. Nature Communications 11:832. doi:10.1038/s41467-020-14644-y

Newlands ES, Stevens MFG, Wedge SR, Wheelhouse RT, Brock C. 1997. Temozolomide: a review of its discovery, chemical properties, pre-clinical development and clinical trials. Cancer Treatment Reviews 23:35–61. doi:10.1016/S0305-7372(97)90019-0

Park D, Morris AR, Battenhouse A, Iyer VR. 2014. Simultaneous mapping of transcript ends at single-nucleotide resolution and identification of widespread promoter-associated non-coding RNA governed by TATA elements. Nucleic Acids Res 42:3736–3749. doi:10.1093/nar/gkt1366

Plosky BS, Frank EG, Berry DA, Vennall GP, McDonald JP, Woodgate R. 2008. Eukaryotic Y-family polymerases bypass a 3-methyl-2’-deoxyadenosine analog in vitro and methyl methanesulfonate-induced DNA damage in vivo. Nucleic Acids Res 36:2152–2162. doi:10.1093/nar/gkn058

Quinlan AR, Hall IM. 2010. BEDTools: a flexible suite of utilities for comparing genomic features. Bioinformatics 26:841–842. doi:10.1093/bioinformatics/btq033

Rhee HS, Pugh BF. 2012. ChIP-exo: A Method to Identify Genomic Location of DNA-binding proteins at Near Single Nucleotide Accuracy. Curr Protoc Mol Biol 0 21 doi:10.1002/0471142727.mb2124s100

Rossi MJ, Kuntala PK, Lai WKM, Yamada N, Badjatia N, Mittal C, Kuzu G, Bocklund K, Farrell NP, Blanda TR, Mairose JD, Basting AV, Mistretta KS, Rocco DJ, Perkinson ES, Kellogg GD, Mahony S, Pugh BF. 2021. A high-resolution protein architecture of the budding yeast genome. Nature 592:309–314. doi:10.1038/s41586-021-03314-8

Sabarinathan R, Mularoni L, Deu-Pons J, Gonzalez-Perez A, López-Bigas N. 2016. Nucleotide excision repair is impaired by binding of transcription factors to DNA. Nature 532:264–267. doi:10.1038/nature17661

Schärer OD. 2013. Nucleotide Excision Repair in Eukaryotes. Cold Spring Harb Perspect Biol 5. doi:10.1101/cshperspect.a012609

Shore D, Nasmyth K. 1987. Purification and cloning of a DNA binding protein from yeast that binds to both silencer and activator elements. Cell 51:721–732. doi:10.1016/0092-8674(87)90095-x

Tubbs A, Nussenzweig A. 2017. Endogenous DNA Damage as a Source of Genomic Instability in Cancer. Cell 168:644–656. doi:10.1016/j.cell.2017.01.002

Wallace SS, Murphy DL, Sweasy JB. 2012. Base Excision Repair and Cancer. Cancer Lett 327:73–89. doi:10.1016/j.canlet.2011.12.038

Weiner A, Hsieh T-HS, Appleboim A, Chen HV, Rahat A, Amit I, Rando OJ, Friedman N. 2015. High-Resolution Chromatin Dynamics during a Yeast Stress Response. Molecular Cell 58:371–386. doi:10.1016/j.molcel.2015.02.002

Whitaker AM, Freudenthal BD. 2018. APE1: A skilled nucleic acid surgeon. DNA Repair (Amst) 71:93–100. doi:10.1016/j.dnarep.2018.08.012

Wyatt MD, Allan JM, Lau AY, Ellenberger TE, Samson LD. 1999. 3-methyladenine DNA glycosylases: structure, function, and biological importance. BioEssays 21:668–676. doi:https://doi.org/10.1002/(SICI)1521-1878(199908)21:8<668::AID-BIES6>3.0.CO;2-D

Yang K, Park D, Tretyakova NY, Greenberg MM. 2018. Histone tails decrease N7-methyl-2′-deoxyguanosine depurination and yield DNA–protein cross-links in nucleosome core particles and cells. PNAS 115:E11212–E11220. doi:10.1073/pnas.1813338115

